# Temporal Dynamics of Neocortical Development in Organotypic Mouse Cultures: A Comprehensive Analysis

**DOI:** 10.1101/2024.04.05.588217

**Authors:** Aniella Bak, Katharina Schmied, Morten Jakob, Francesco Bedogni, Olivia Squire, Birgit Gittel, Maik Jesinghausen, Kerstin Schünemann, Yvonne Weber, Björn Kampa, Karen M. J. van Loo, Henner Koch

## Abstract

Murine organotypic brain slice cultures have been widely used in neuroscientific research and are offering the opportunity to study neuronal function under normal and disease conditions. Despite the brought application, the mechanisms governing the maturation of immature cortical circuits *in vitro* are not well understood. In this study, we present a detailed investigation into the development of the neocortex *in vitro*. Utilizing a holistic approach, we studied organotypic whole-hemisphere brain slice cultures from postnatal mice and tracked the development of the somatosensory area over a five-week period. Our analysis revealed the maturation of passive and active intrinsic properties of pyramidal cells together with their morphology, closely resembling *in vivo* development. Detailed Multi-electrode array (MEA) electrophysiological assessments and RNA expression profiling demonstrated stable network properties by two weeks in culture, followed by the transition of spontaneous activity towards more complex patterns including high-frequency oscillations. However, weeks 4 and 5 exhibited increased variability and initial signs of neuronal loss, highlighting the importance of considering developmental stages in experimental design. This comprehensive characterization is vital for understanding the temporal dynamics of the neocortical development *in vitro*, with implications for neuroscientific research methodologies, particularly in the investigation of diseases such as epilepsy and other neurodevelopmental disorders.

## Introduction

Due to the limitations of experimental animal models, which are costs, labor and ethical concerns on animal numbers and suffering, many advances have been made during recent years to improve *in vitro* neurological models to better mimic the *in vivo* situation. Although induced pluripotent stem cells (iPSCs) and organoids have been a big step in the right direction and provide the advantage of recapitulating human development in simplified neural networks^1^, partially preserve genetic patient characteristics^2^ and eliminate the ethical concerns tied to the use of embryonic stem cells^3^, they unfortunately still encounter the problem of impaired maturation and lack of cellular diversity and structural complexity^4,5^. While acute brain slices are a viable option to investigate the rodent brain at different ages relatively close to the *in vivo* situation, they unfortunately do not enable genetic modification, long-term stimulation or pharmacological manipulation. Although these methods can be applied in the living animal, *in vitro* preparations offer more precisely controlled experimental conditions. Organotypic brain slice cultures (oBSCs) close this gap by providing an experimental time frame of multiple weeks along with intact three-dimensional cytoarchitecture as well as inclusion of all cellular subtypes of the brain. oBSCs have been widely adapted in neuroscientific research since they were first established by Gähwiler^6^ with the initial roller tube culturing technique. Though this method is robust and yields stable cultures for multiple weeks^7–11^, the thinning to 2-3 cell layers and therefore loss of the original spatial organization might be preferable for applications where visual accessibility is favored, but not for electrophysiological, morphological and developmental studies that require intact three-dimensional architecture of certain brain areas. As a consequence, the method was further adapted by Stoppini et al.^12^, introducing a semiporous membrane culturing method at air-liquid-interface to preserve the original structure of brain preparations. Both methods have been shown to maintain the organotypic structure, their normal cell morphology and to generally correlate well with development observed *in vivo*^7,9,13–17^ and have been used successfully in the past, investigating, for example, neurodegeneration^18,19^, ischemia^20–23^, interneuron development^7,10^, axon formation and dynamics^24,25^ and network dynamics^15,26,27^. In addition, oBSCs have been used as a screening platform for novel therapy targets and therapeutics^14^.

Especially preparations of the rat and mouse hippocampus have been used and characterized widely ^11,28–33^ with only a limited number of studies reporting the use of other brain areas such as cerebellum^13,15,34,35^, striatum^10,36^ or cortex^7,9,16,37–39^. Neocortical oBSCs are of special interest when it comes to the investigation of inhibition and excitation balancing, developmental network dynamics, cell-cell relationships as well as pathogenesis in the maturing cortex. Unfortunately, to date, information on the tissue development *ex vivo* is scarce and has only been reported in cultures created by the initial roller tube technique^7–10,25,40–42^ or in studies only focusing on specific parameters^24,37–39,43^. This study aimed to close this gap of knowledge by providing a holistic characterization of the development of membrane insert cultured mouse neocortical oBSCs over a time course of five weeks, investigating single cell and network electrophysiology, RNA expression levels and morphology of pyramidal cortical neurons, as this information will hold vital indications on which timeframes of culturing are suitable for different types of experimental approaches.

## Material & Methods

### Animals

All procedures involving animals were performed in accordance with the guidelines of the Federation of European Laboratory Animal Science Association, the EU Directive 2010/63/EU, and the German animal welfare law. For harvesting of brain tissue for organotypic cultures, wildtype C57BL/6J mice of either sex aged 5-7 postnatal days were used. Mice were obtained from the local Institute of Laboratory Animal Science RWTH University Clinic Aachen and housed at a 12h light-dark cycle with food and water ad libitum. Tissue harvest was performed after decapitation of mice.

### Slice preparation

After mouse decapitation, the brain was carefully extracted from the skull using microdissection tools. Hemispheres were separated and glued on solidified agarose blocks with rostral side facing upwards. Brain tissue was sliced at 350 µm and whole-hemisphere slices were used for cultivation. Brain slices were generated using a vibratome (Leica VT1200S) filled with ice-cold artificial cerebrospinal fluid (aCSF), containing: 125 mM NaCl, 25 mM NaHCO_3_, 2.5 mM KCl, 1.25 NaH_2_PO_4_, 1 mM MgCl_2_ * 6 H_2_O, 2 mM CaCl_2_, 25 mM glucose and 1% penicillin/streptomycin (Gibco) dissolved in Millipore water. aCSF was constantly perfused with carbogen gas (95% O_2_ and 5% CO_2_) and kept cold by reusable ice cubes for appropriate hypothermia and oxygenation during preparation. For all experiments except RT-PCR, somatosensory cortical regions of whole mouse brain slices were used and analyzed.

### Organotypic brain slice cultures

Slices were placed on semipermeable membrane inserts (pore size 0.4 µm, Millipore), adhered by removing excess liquid and placed in 6-well plates at air-liquid interface on 1.2 ml of controlled medium containing: 20% horse serum, 1 mM L-glutamine, 0.00125% ascorbic acid, 0.001 mg/ml insulin, 1 mM CaCl_2_, 2 mM MgSO_4_, 13 mM glucose and 1% penicillin/streptomycin in minimum essential medium at an osmolarity of 320 mOsm/l and pH of 7.28. Cultures were maintained in an incubator with controlled atmosphere (37°C, 5% CO_2_, 100% humidity) and full medium change was performed every two days.

### Time periods

All experiments on organotypic brain slice cultures were performed in a time frame of 1-37 days *in vitro*. For Patch-Clamp experiments, periods are defined as initial (1-2 DIV), week 1 (6-9 DIV), week 2 (11-16 DIV), week 3 (17-23 DIV), week 4 (28-30 DIV) and week 5 (34-37 DIV). Remaining experiments were performed on the exact corresponding day, meaning initial (2 DIV), week 1 (7 DIV), week 2 (14 DIV), week 3 (21 DIV), week 4 (28DIV) and week 5 (35 DIV), +/- 1 DIV.

### Patch-Clamp recordings

Whole-cell Patch-Clamp recordings on mouse organotypic brain slice cultures were performed at various time points with cultures being *in vitro* between 2 – 37 days. For this cause, brain slices were excised from the cultivation membrane and transferred to the recording bath. During recordings, slices were continuously perfused with carbogenated aCSF at 30°C at a perfusion rate of approximately 7 ml/min. Recording pipettes were pulled from borosilicate glass capillaries (World Precision Instruments) with a resistance of 3-7 MΩ. For recordings, pipettes were filled with intracellular solution containing: 140 mM K-gluconic acid, 1 mM CaCl_2_, 10 mM EGTA, 2 mM MgCl_2_, 4 mM Na_2_ATP, 10 mM HEPES and 5 mg/ml biocytin (Sigma). Neurons were patched within the somatosensory cortical region of whole mouse brain slices, approximately between 100 – 500 µm from the cortical border. Pyramidal cells were identified by their somatic shape and firing properties. Whole-cell recordings were performed using an EPC10 amplifier (HEKA) and Patchmaster software (HEKA). Signals were sampled at 10 kHz, filtered at 1 kHz and analyzed using Fitmaster software (HEKA) and custom written MatLab (MathWorks) scripts. After recordings, slices were fixed in 4% PFA (Morphisto) for 10-24 h.

### Multi-electrode array recordings

Multi-electrode array (MEA) recordings on mouse organotypic brain slice cultures were performed at time points 2 DIV, 7 DIV, 14 DIV, 21 DIV, 28 DIV and 35 DIV. Brain slices were excised from the cultivation membrane and transferred onto the MEA chip (256MEA30/8iR-ITO-pr, 16×5616 electrodes, electrode size 30 μm, 200 μm spacing, Multi Channel Systems MCS GmbH) with the electrode grid covering most of the slice, especially the somatosensory cortical region. Slices were weight down by a meshed harp and continuously perfused with carbogenated aCSF at a temperature of 30°C. Recordings were performed using a 256 electrode MEA system (USB-MEA 256, Multi Channel Systems MCS GmbH) with an electrode size of 30 µm and in-between electrodes spacing of 200 µm. Signals were sampled at 10 kHz using Multi Channel Experimenter software (Multi Channel Systems MCS GmbH). After an equilibration period of 20 min, slices were recorded for 2 x 5 min and recordings were analyzed using custom scripts in MatLab (MathWorks).

### Parvalbumin staining

Free-floating organotypic slices were washed in PBS-T (0.1% Tween, Invitrogen; PBS 1x) for 10 minutes and then incubated in 0.1M glycine in PBS for 30 minutes. Slices were washed in PBS-T (3 times, 15 minutes each) and incubated in blocking solution (10% horse serum, 1% triton in PBS) for 1 hour, after which sections were incubated at 4°C with NeuN (1:500; Millipore, ABN78) and parvalbumin (1:1000; Sigma, P3088) antibodies for 48 hours. Following this, slices were thoroughly washed in PBS-T (10 times, 15 minutes each), and then incubated in the secondary antibody mix, consisting of AlexaFlour secondary antibodies (1:500, ThermoFisher) and DAPI (1:500) in blocking solution, for 1 hour at room temperature. Slices were then washed twice more in PBS-T, and mounted on slides with Fluoromount (Invitrogen). Images were acquired using either a Leica Upright Microscope or Zeiss LSM710 laser scanning confocal microscope.

### RNA isolation and cDNA generation

For RNA isolation, two whole hemisphere slices from the same culturing insert were collected into one tube to obtain sufficient amounts of RNA. Tissue was homogenized by addition of one metal bead (steel, Qiagen) and 1 ml trizole (Invitrogen) per tube and inserting them into a SpeedMill (Analytik Jena) for 2 minutes. Subsequently, samples were centrifuged for 1 min at 1000 rpm and 4°C, left on ice for 30 min, homogenized and then left at room temperature for 5 min. 200 µl of Chloroform/Trichloromethane (Sigma) were added to each tube, vortexed for 60 seconds and left at room temperature for 3 min. Samples were then centrifuged at 12.000 g and 4°C for 30 min and aqueous phase was pipetted into new tubes. 400 µl isopropanol was added to each tube, mixed and left at room temperature for 10 minutes. Subsequently, samples were centrifuged for 10 min at 12.000 g and 4°C, supernatant was discarded and 800 µl 75% ethanol were added. This step was repeated for a second time, then tubes were left upside down on a clean tissue to dry at room temperature. The pellet was finally taken up in 25 µl nuclease free water each and the amount of RNA was measured on a NanoDrop.

All samples showing sufficient RNA concentration (> 100 ng/ul) and purity (260/280 ratio of 2.0 +/- 0.2) were then used to generate cDNA by normalizing the sample with the lowest concentration to 15 µl volume and calculation of the amount of RNA for remaining samples accordingly. To each sample RNA, 4 µl of iScript reaction mix (Bio-Rad), 1 µl of iScript reverse transcriptase (Bio-Rad) and nuclease free water to fill total volume to 20 µl were added. cDNA was finally generated in a Thermocycler (Biometra) by priming at 25°C for 5 min, reverse transcription at 46°C for 20 min and reverse transcriptase inactivation step at 95° for 1 min.

### RT-PCR

mRNA quantification was performed by real-time RT-PCR using the ΔΔCt-method. Quantitative PCR was performed in an QuantStudio^TM^ I apparatus (Applied Biosystems) containing 1x iQ SYBR Green supermix (Bio-Rad Laboratories Inc.), 5 pm of each oligonucleotide primer (*Actb*: 5ʹ-CATTGCTGACAGGATGCAGAAGG-3ʹ and 5ʹ-TGCTGGAAGGTGGACAGTGAGG-3ʹ; *Syp*: 5ʹ-TTCAGGACTCAACACCTCGGT-3 and 5ʹ-CACGAACCATAGGTTGCCAAC-3ʹ; *Scn1a*: 5ʹ-CTTGAGCCCGAAGCTTGCT-3ʹ and 5ʹ-TCCTTCTTCCACGCTGATTTG-3ʹ; *Scn2a*: 5ʹ-CATCGTCTTCATGATTCTGCTCA-3ʹ and 5ʹ-GGTTTTTCGCTGCTCGATGTA-3ʹ; *Scn8a*: 5ʹ-CATCTTTGACTTTGTGGTGGTCAT-3ʹ and 5ʹ-TGACGCGGAATAGGGTCG-3ʹ; *Pvalb*: 5ʹ-CTGGGGTCCATTCTGAAGGG-3ʹ and 5ʹ-TTCAACCCCAATCTTGCCGT-3ʹ; *Hcn1*: 5ʹ-TGAAGCTGACAGATGGCTCTT-3ʹ and 5ʹ-GAGTCGGTCAATAGCAACTGTCT-3ʹ) and 2.5-fold diluted synthesized cDNA (total volume of qPCR reaction 20 μl), by incubating 3 min at 95°C, 40 cycles of 15s at 95°C and 1 min at 60°C. Quantification was based on β-actin.

### Biocytin stainings

Cells previously filled with biocytin during Patch-Clamp recordings were washed three times for 20 minutes in Phosphate buffered saline (PBS-T-T) containing 137 mM NaCl, 2.7 mM KCl, 10 mM Na_2_HPO_4_, 1.8 mM KH_2_PO_4_, 0.05 % Triton X-100 (Sigma) and 0.2 % Tween-20 (Sigma-Aldrich). Subsequently, slices were incubated in a staining solution containing 0.01 mg/ml Streptavidin-Cy3 (Sigma-Aldrich) in PBS-T-T overnight at 4°C. On the following day, slices were washed three times with PBS for 10 minutes, incubated in DAPI (Sigma, 1.43 µM) solution for 5 minutes and again washed three times with PBS for 10 minutes. Finally, slices were mounted on Superfrost Plus adhesion slides (Epredia) with Fluoromount G (ThermoFisher Scientific).

### Confocal imaging and morphological reconstructions

Neurons stained with Streptavidin-Cy3 were selected for confocal imaging if they exhibited sufficient fluorescent signal and overall morphology of a pyramidal cell. Cells were then scanned as a whole in x, y and z axis on an inverted confocal microscope (Zeiss LSM710 and Zeiss LSM980 Airyscan 2) with a 40x oil immersion objective. Single stacks were imaged at a resolution of 1024×1024 with multiple tiles in x and y direction and z-stacks at 0.66 μm thickness. Fluorophore Cy3 was imaged with a 561 nm laser and a 569-712 filter.

Using the complete confocal neuron scans, 3D models of corresponding neurons were generated using the neuron tracing tool in Arivis software (Zeiss). Axon, dendrites and soma were segmented and obtained 3D models of neurons were further used for morphological analysis.

### Patch-Clamp recording analysis

#### Data analysis and statistics

Patch-Clamp data was analyzed using Fitmaster software (HEKA) and custom written scripts for MatLab (MathWorks). Cells were included in analysis, when membrane potential was <= - 60 mV, they were able to generate multiple action potentials upon current injection and action potential amplitude was larger than 40 mV. In Fitmaster, data was filtered digitally at 1 kHz to remove artefacts. Action potentials were analyzed using the corresponding software function. The membrane potential was measured at the start points of every protocol and a mean value was calculated for every cell. Input resistance was determined by creating an I-V curve of current input responses in whole-cell mode and calculating the reverse slope value delta V/I. Spontaneous events in current-clamp mode were characterized as starting by reaching rheobase (at least one spike) and ending if membrane potential returned to baseline for at least 500 ms. Spontaneous events in voltage clamp mode were characterized by reaching an amplitude of at least 200 pA and ending if current returned to baseline for at least 500 ms. Spontaneous events were analyzed using the corresponding functions in Fitmaster software. Several integral values that could not be calculated in Fitmaster due to baseline shift issues were subsequently calculated with a custom-written MatLab script. For all values corresponding to a single event (duration, integral, spikes, pos./neg. amplitude), mean values were created for every cell. If Gaussian distribution was assumed (membrane potential, input resistance, AP Half-width, voltage sag), ROUT method outlier test was performed and significance in differences between time periods was calculated per GraphPad Prism 10 software using One-Way ANOVA or mixed statistical test with post-hoc Tukey’s multiple comparisons test. Remaining data was assumed to be non-parametrical and therefore analyzed with Kruskal-Wallis statistical test and Dunn’s multiple comparison correction.

#### MEA analysis

MEA recordings were analyzed using custom-written Matlab (MathWorks) scripts. For each slice, two recording epochs of 5 minutes and (if present) three network-driven local field potentials (LFPs) per recording were analyzed. The data was first filtered with a low pass filter of 50 Hz and, if present, noisy channels were removed from the analysis. LFP events were detected at individually selected threshold per recording in a manually chosen reference channel. An individual time window was subsequently set for the analysis of an LFP event. In a first step, we qualitatively judged the complexity of the LFP as a) very low b) low c) medium d) complex or d) very complex. Categorization criteria were as follows: one single spike (very low), 2-3 spikes (low), >3 spikes (medium), initial spike followed by high-frequency oscillation at a low amplitude not reaching the initial amplitude (complex), high-frequency oscillation at a high amplitude reaching the full initial amplitude (very complex) or not applicable (n. a.), when the waveform did not fall into one of the categories. Subsequently, the average of the minimum and maximum amplitudes over all channels over the duration of the recording was calculated. The threshold was set for positive and negative LFP components as a value between 5-20% above or below the average value. Afterwards, the electrodes that crossed the threshold values at least once within the selected recording period were counted as active electrodes. From the number of active electrodes we further calculated 1) the area on the electrode grid covered by LFP activity, 2) a ratio of positive vs. negative electrodes (note that an electrode could be both positive and negative if the signal crossed both thresholds during the period analyzed), 3) the absolute summation of LFP amplitudes (cumulative most negative and positive values) and average LFP amplitude (averaged over all active electrodes) of the positive and negative LFP, 4) duration of the LFP (averaged over all channels). Significance in differences between time periods was calculated using Kruskal-Wallis statistical test with post-hoc Dunn’s multiple comparisons test per GraphPad Prism 10 software.

#### Staining analysis (Parvalbumin)

Per slice, two regions of interest were selected, one in the prefrontal cortex/secondary motor cortex and one in somatosensory cortex. As no difference was found between those two regions, both ROIs were counted and included in analysis. Cell counting was automated and NeuN count was used as normalizer at 100% to calculate overall percentage of Parvalbumin-positive cells. Mean values were generated for every slice and subsequently, for every time period. Significance in differences between time periods was calculated using One-Way ANOVA statistical test with post-hoc Tukey’s multiple comparisons test per GraphPad Prism 10 software.

#### Morphological analysis

Neurons were reconstructed with the neuron tracing tool in Arivis software (Zeiss). Dendrites were reconstructed from the soma to terminal end points and 3D models of neurons were subsequently segmented into apical and basal dendrites. Values for dendritic length, branch points and terminal end points were ascertained in Arivis software and plotted in GraphPad Prism 10 software. The axon was not included in the analysis. Significance in differences between time periods was calculated using One-Way ANOVA statistical test with post-hoc Tukey’s multiple comparisons test.

## Results

### Passive and firing properties

For this study, we used organotypic brain slice cultures (oBSCs) generated from P5-6 mice and targeted pyramidal cells of the somatosensory area of the cortex, located at a depth of 100 – 500 μm from the cortical surface. Pyramidal cells were identified during Patch-Clamp recordings by their somatic shape and general firing behavior, as well as post-hoc by their morphology (figure 2A). Recordings were performed continuously from 2 days *in vitro* (DIV) up to 37 days *in vitro* with six time periods characterized as: initial (1-2 DIV), week 1 (6-9 DIV), week 2 (11-16 DIV), week 3 (17-23 DIV), week 4 (28-30 DIV) and week 5 (34-37 DIV). Overall, 179 neurons were recorded and included in our analysis for time periods initial (n=21), week 1 (n=28), week 2 (n=36), week 3 (n=45), week 4 (n=30) and week 5 (n=19). In response to depolarizing current injection, we observed that neurons of the initial culturing period exhibited immature firing behavior and depolarization block after several action potentials, thus not being able to fire continuous trains of action potentials when depolarized into firing saturation (figure 2B, initial). Interestingly, this observation resolved already during the first week in culture with neurons gaining and retaining the ability to fire continuous trains of action potentials up to 5 weeks in culture (figure 2B, week 1 – week 5). While nearly two thirds of recorded neurons displayed a depolarization block during the initial culturing period, this phenomenon was only found occasionally during the later timepoints (figure 2C). Action potentials were broad during the initial culturing period and the first week with an action potential half-width of 2.62 ± 0,15 ms and 2.44 ± 0,11 ms, respectively and decreased continuously during the following weeks to reach mature steady-state values of 1,86 ± 0,10 ms after 3 weeks (figure 2D). The instantaneous firing frequency was elevated during the initial culturing period with an area under the curve (AUC) value of 269,3 ± 31,04 but decreased and stabilized during culturing weeks 1 and 2 to a value of 124,5 ± 16,07. Interestingly, this value increased again at the later culturing periods to a final value of 528,8 ± 65,28 in week 5 (figure 2E).

**Figure 1.**
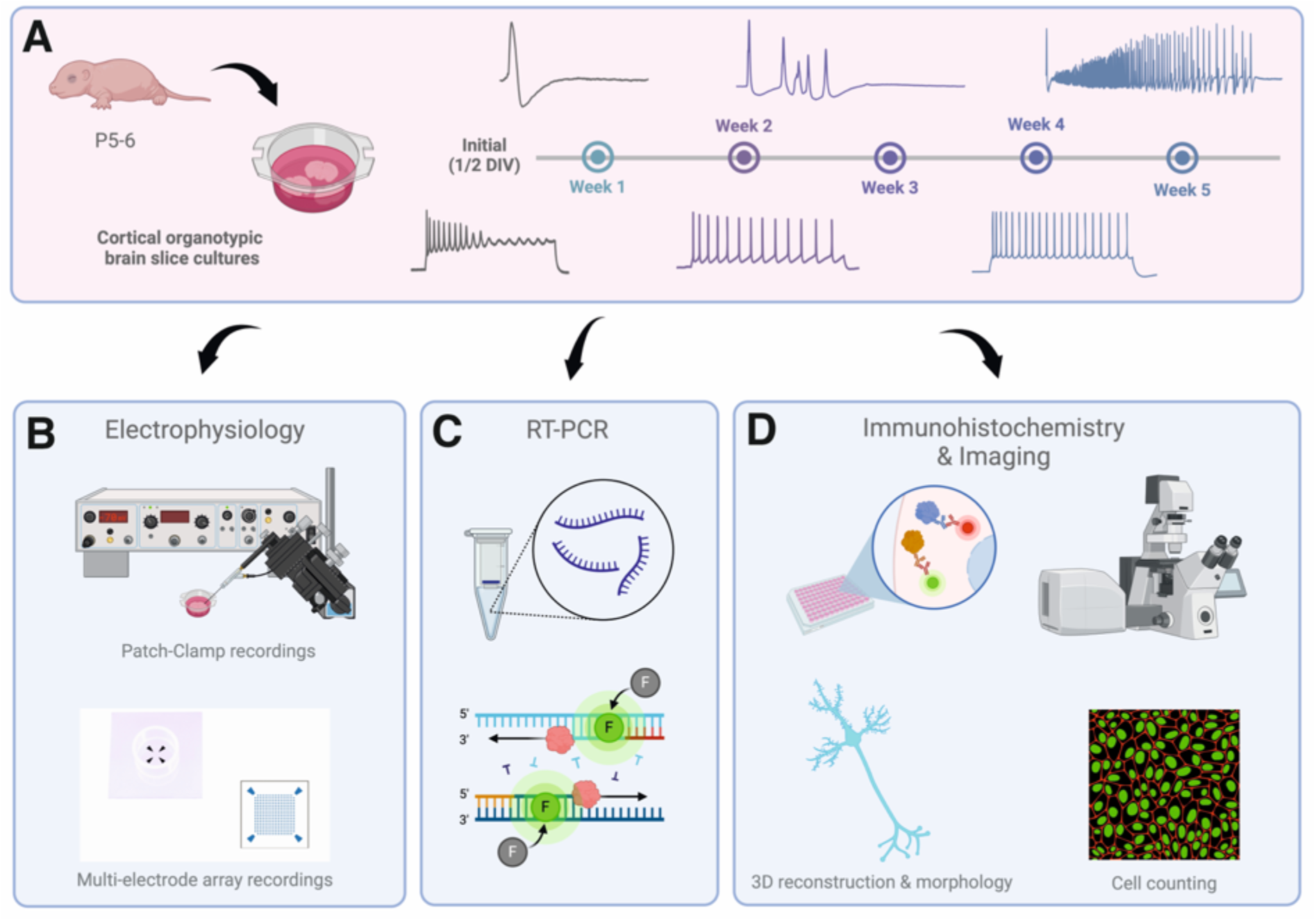
Graphical abstract. Schematic overview of experimental setup. (A) Cortical organotypic brain slice cultures were created from P5-6 wildtype mice and cultured on semiporous membrane inserts in controlled medium. Experiments were performed continuously over a time course of 5 weeks. Development of single-cell firing properties and network events displayed along the timeline. (B) Electrophysiological data was obtained by performing whole-cell Patch-Clamp recordings and Multi-electrode array (MEA) recordings. (C) Real-time PCR was performed on cDNA samples created from isolated RNA from cultures to monitor RNA expression levels. (D) Native cultures were stained for cellular expression markers and neurons filled during Patch-Clamp recordings were stained and reconstructed for morphological analysis.

**Figure 2.**
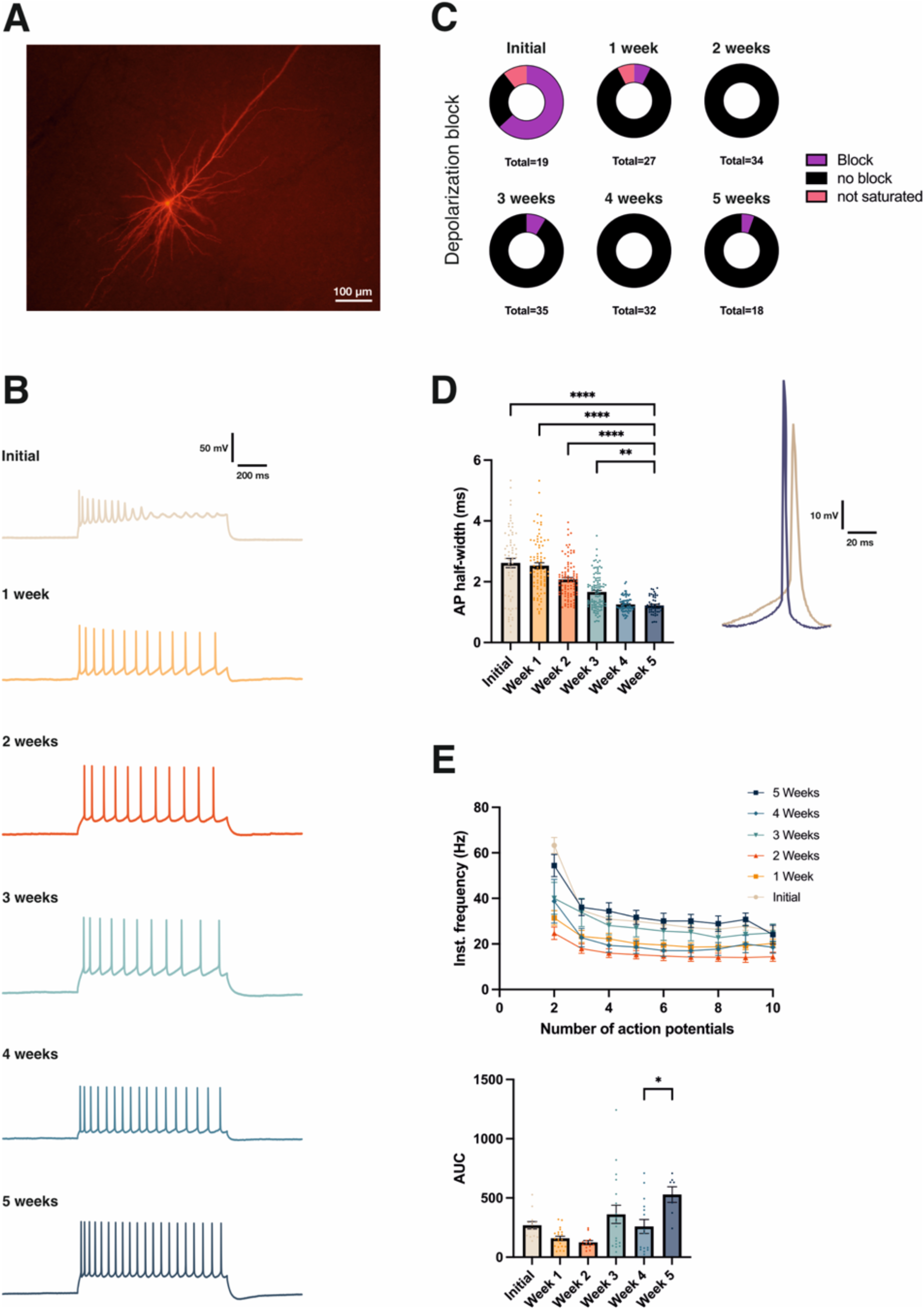
Firing properties of recorded neurons. (A) Exemplary pyramidal cell, biocytin filled during Patch-Clamp recordings and stained with Streptavidin-Cy3, 16 DIV. (B) Examples of induced spike trains from pyramidal cells of every time period, approximately 100 pA above rheobase. (C) Illustration of proportion of neurons exhibiting depolarization block upon current injection until saturation of number of action potentials. (D) Development of action potential half-width over a time course of five weeks, **** = p<0.0001 for One-way ANOVA test. Visual comparison of action potential width between initial culturing period and five weeks *in vitro*. (E)Development of instantaneous firing frequency over five weeks of cultivation, area under the curve illustrated in the graph below, * = p<0.05 for Kruskal-Wallis test.

Another parameter for neuronal maturation in organotypic cultures was found in the voltage sag, defined as the ratio between the steady-state decrease and the largest decrease in voltage following a hyperpolarizing current, in this case - 200 pA. The sag is caused by hyperpolarization-activated cation current (*I*_h_) and thought to be mainly controlled by activation of HCN channels^44^. The voltage sag was significantly elevated during the initial culturing at 9,07 ± 1,04 mV, but decreased significantly and reached physiological steady-state levels at week 1 with 3,14 ± 0,47 mV (figures 3A and 3C). The cellular input resistance displayed a similar development over time with values elevated during initial culturing at 338,2 ± 29,19 MΩ, followed by a significant decrease and stabilization after one week to 168 ± 9,91 MΩ (figures 3B and 3C). The resting membrane potential resided at - 73,65 ± 0,96 mV in the initial culturing phase and underwent significant hyperpolarization by week 2, dropping to - 81,19 ± 1,72 mV (figure 3D). All individual values are given in supplementary table 1.

**Figure 3.**
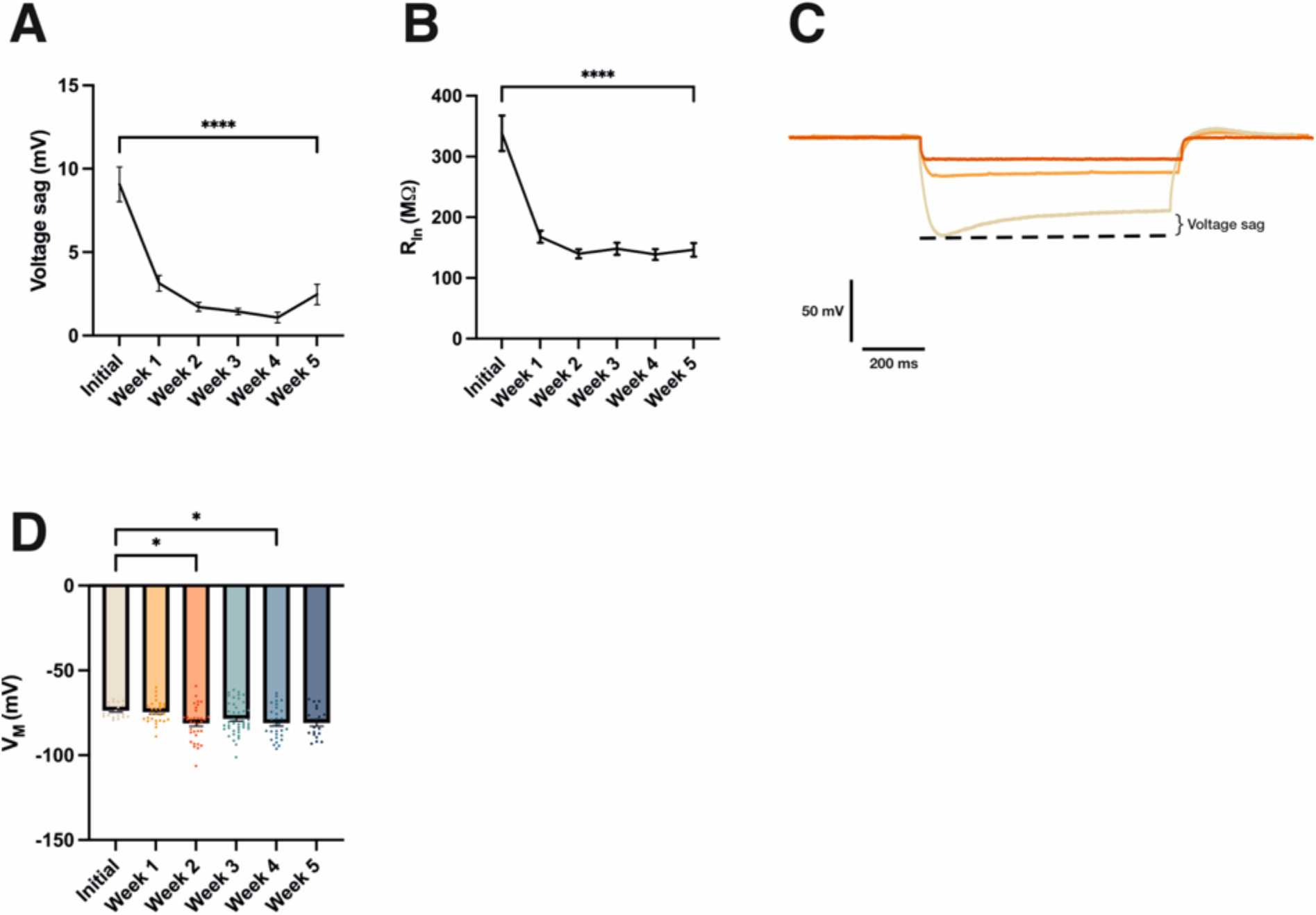
Passive proper’es and voltage sag of recorded neurons. (A) Development of voltage sag over five weeks. (B) Development of input resistance over five weeks. (C) Comparison of input resistance and voltage sag between neurons of ini9al, week 1 and week 2 culturing periods. (D) Development of res9ng membrane poten9al over a 9me course of five weeks. Significance levels for all graphs: * = p<0.05, **** = p<0.0001 for One-way ANOVA test.

Overall, our data suggested a generally immature neuronal phenotype shortly after culture production from mice aged 5-7 postnatal days, but also continuing maturation during the first three weeks *in vitro* to a more stabilized state with increasingly physiological firing behavior and intrinsic excitability.

### Development of spontaneous activity in organotypic neocortical cultures

We first wanted to observe spontaneous activity in organotypic cultures on a single-cell level and therefore monitored events in current-clamp mode during whole-cell Patch-Clamp recordings over our culturing periods initial (n=17), week 1 (n=25), week 2 (n=30), week 3 (n=36), week 4 (n=23) and week 5 (n=17). This data set contained less cells than the previous one, as recordings of spontaneous activity were not performed for every cell. Spontaneous activity in current-clamp mode was characterized as an “event” when it reached action potential threshold at least once and was considered as finished when it returned to baseline afterwards for at least 500 ms (figure 4A).

**Figure 4.**
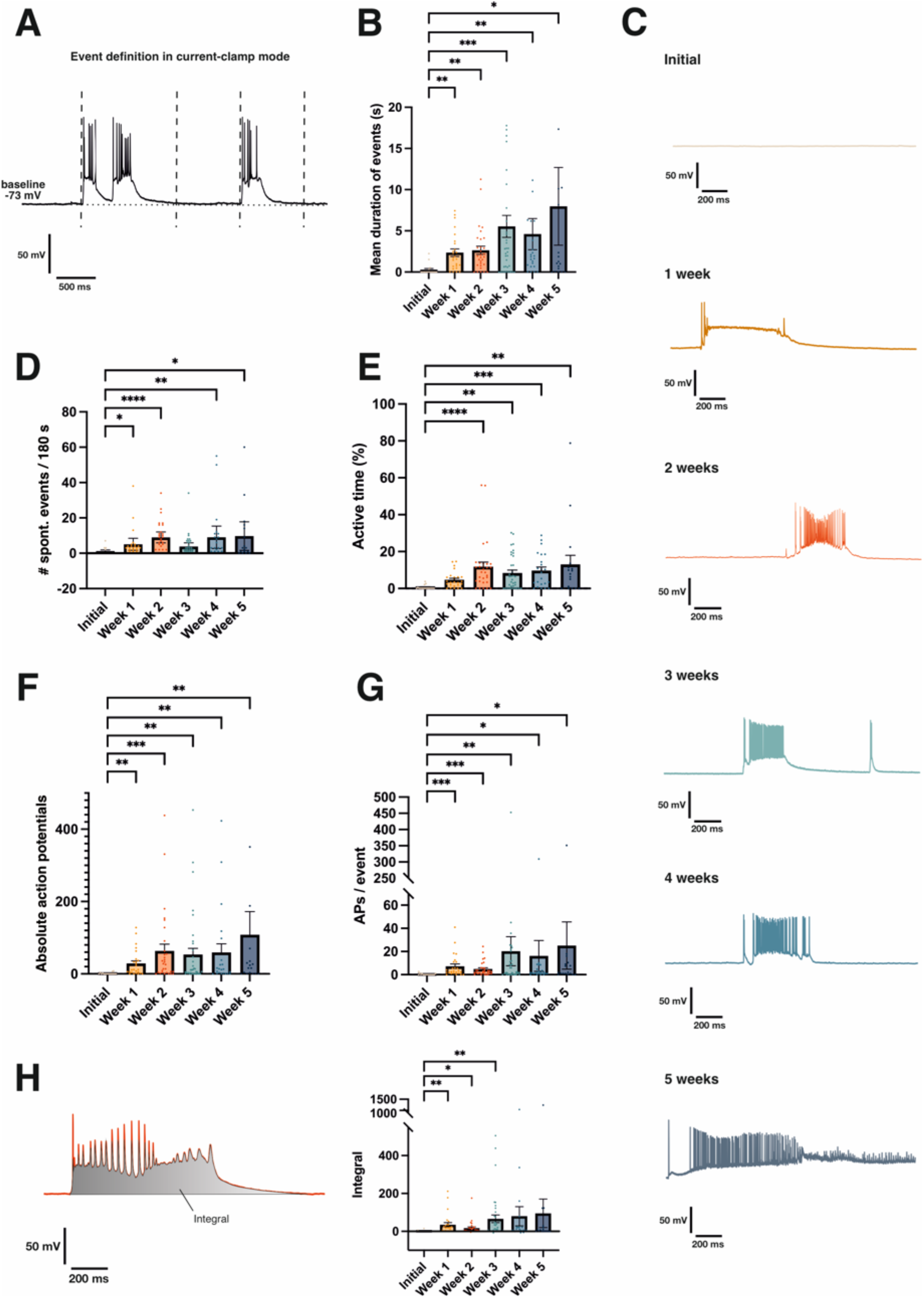
Spontaneous activity as recorded in current-clamp mode. (A) Definition of events during current-clamp recordings, event starting at reaching action potential threshold and ending when returned to baseline for at least 500 ms. (B) Mean duration of spontaneous events. (C) Exemplary traces of spontaneous up-states recorded. (D) Number of spontaneous events recorded per cell over a recording time of 3 minutes. (E) Percentage of recording cell spent in active state, meaning above baseline in active event as defined in A. (F) Overall number of spontaneous action potentials recorded in one cell over a recording time of 3 minutes. (G) Number of action potentials per single event recorded. (H) Visual representation of definition of integral as area under the curve of a spontaneous event. Graph illustrating numeric evaluation depicted on the right. Significance levels for all graphs: **** = p<0.0001, *** = p<0.001, ** = p<0.01, * = p<0.05 for Kruskal-Wallis-Test.

We evaluated a recording time of three minutes for every cell and calculated mean values for every parameter corresponding to single events, such as duration, action potentials per event and integral. We observed a striking increase in number of events and event duration over the cultivation period of five weeks, with neurons exhibiting nearly no spontaneous activity during the initial culturing phase and becoming gradually more active over time. The mean duration of spontaneous events increased from short bouts of activity to more complex and elaborate activity patterns which still contained single spikes and short events, but also cellular up-states. This resulted in a significant increase of the mean duration of events between the initial culturing period, where events displayed a mean duration of 0,27 ± 0,16 s, and following time periods, with a mean duration of 2,37 ± 0,43 s at week 2 and 7,97 ± 4,72 s at week 5 (figure 4B). The development of spontaneous activity is best illustrated by the exemplary traces shown (figure 4C), showing the mostly silent state of initial cultures (figure 4C, initial) progressing to up-states exhibiting only few action potentials and partial depolarization block (figure 4C, week 1) to elaborate spontaneous events accompanied by prolonged bursts of action potentials during later cultivation periods (figure 4C, weeks 2, 3 and 4). Interestingly, several neurons started displaying abnormal forms of excitatory discharges at week 5 in culture (figure 4C, week 5), however, it has to be stressed that these high frequency events were not the exclusive or dominating form of spontaneous activity and were only rarely observed in cultures aged 4-5 weeks (supplement figure 1). This is also highlighted by the fact that spontaneous activity did not seem to continuously accelerate into a high-frequency phenotype during culture as shown by the overall number of events per cell, with on average 0,8824 ± 0,46 events observed per 180 s in the initial culturing period and stable values of 8,967 ± 1,52 events by week 2 (figure 4D). This was also reflected by the active time spent above resting membrane potential, increasing from 0,51 ± 0,27% at initial culturing periods to a steady-state value of 11,81 ± 2,48% by week 2 (figure 4E). These activity parameters only showed a significant increase when comparing the initial culturing phase to later culturing periods, but no continuous increase in-between later periods. Instead, spontaneous activity shifted to more complex patterns composed of single spikes, bursts and long-lasting up-states. The absolute action potentials recorded per cell significantly increased after the initial culturing from 1,235 ± 0,63 to 63,7 ± 18,58 at week 2, stayed stable until week 4 and subsequently showed a trend towards increasing further in week 5 to 108,2 ± 64,22 (figure 4F). The change between the proportions of different types of spontaneous activity was especially highlighted by the number of spikes per event, which saw a significant increase from initial values at 0,35 ± 0,18 to 4,88 ± 1,10 at week 2, but also a clear trend in late time periods towards spontaneous events encompassing larger numbers of action potentials with a final value of 25,14 ± 20,41 at week 5 (figure 4G). This was further illustrated by calculating the integral (the area under the curve) for every event and comparing mean values between individual cells. The integral value increased significantly from 1,23 ± 0,93 Vs at the initial culturing phase compared to 66,19 ± 20,38 Vs at week 3, but the difference ceased to be significant in week 4 and 5 despite values of 79,85 ± 50,43 Vs and 95,11 ± 75,48 Vs, respectively. This was due to the activity of the neurons becoming more heterogenous, resulting in a rise of variance and a clustering into cells exhibiting predominantly single spikes and short bursts versus cells that tended to display long, elaborate up-states, some of them potentially drifting into either high-frequency oscillations or runaway excitation and pathological activity. All individual values are given in supplementary table 1.

In summary, when looking at the intracellular spontaneous activity recorded at the plasma membrane of the cell, cortical pyramidal neurons were near silent in the initial phase of cultivation, but started to progressively develop physiological spontaneous activity during the first two weeks in culture (figure 5, week 1). For later cultivation time points, especially week 4 and 5, heterogeneity between individual cells as well as cultures rose, including all kinds of spontaneous activity patterns from near-silent, single spiking and bursting activity to complex up-states and even discharge patterns that could be characterized as high-frequency oscillations or epileptiform activity (figure 5, week 4).

**Figure 5.**
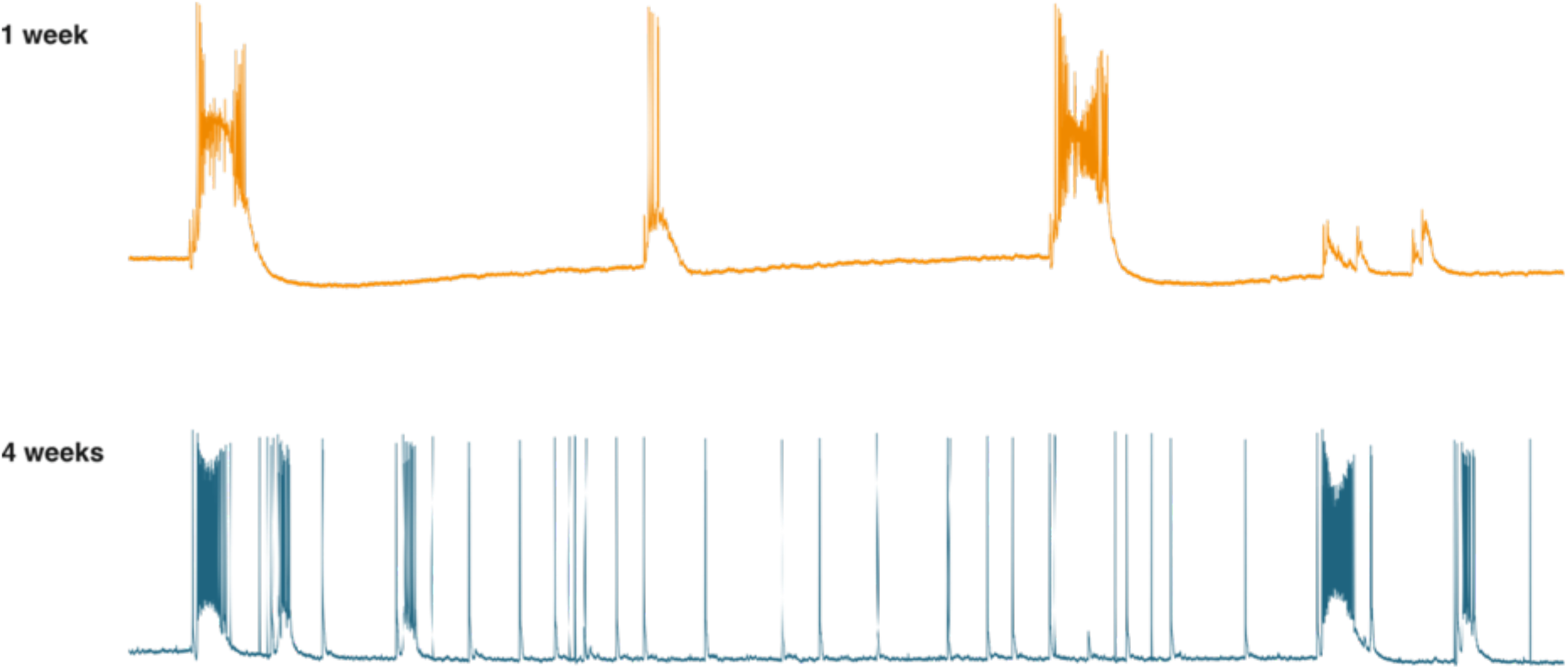
Spontaneous network activity of organotypic cortical cultures after 1 week vs. 4 weeks. Depiction of three minutes of current-clamp whole-cell Patch-Clamp recordings of pyramidal neurons in mouse cortex during week 1 vs. week 4 in culture. Up-states were found in both periods, however, up-states found during week 1 of culturing were lower in frequency and displayed partial depolarization block, which was resolved in later culturing periods.

To get a better idea of how these patterns of spontaneous activity develop, we next analyzed voltage-clamp recordings of whole-cell Patch-Clamp and therefore the synaptic input individual cells were receiving. Again, we included cells over the whole time course of our culturing periods, namely initial (n=14), week 1 (n=17), week 2 (n=21), week 3 (n=34), week 4 (n=23) and week 5 (n=17). Spontaneous activity in voltage-clamp mode was characterized as an “event” when it reached a threshold of at least 200 pA and considered as finished when it returned to baseline afterwards for at least 500 ms. Events were recorded at a holding potential of -70 mV and negative as well as positive amplitudes were calculated from the baseline (figure 6A). As expected, similar to the events recorded in current-clamp, the number of spontaneous synaptic events increased significantly from 1.0 ± 0,46 at the initial culturing period to a steady value of 11,1 ± 1,39 at week 2 (figure 6B), but rose again in week 5 to 16,12 ± 6,15. Also matching the trend seen in current-clamp recordings was the mean duration of events occurring in voltage-clamp mode, which increased significantly from 0,32 ± 0,14 s at the initial culturing period to 3,40 ± 0,90 s in week 1 and 8,57 ± 2,48 s in week 4 (figure 6C). Interestingly, we found a trend towards decreasing mean event duration in week 5 to 4,4 ± 1,56 s, which is in contrast to the increase observed in current-clamp recordings. Regarding the mean negative amplitude of synaptic events, which reflected the excitatory input the neuron was receiving, we observed a significant increase from -267,9 ± 193,2 pA at the initial culturing period to -1337 ± 324,8 pA in week 1, followed by a continuous decrease of amplitude with the decline being significant between week 1 and week 5 with a final value of -394,9 ± 101,8 pA (figure 6D). This result might seem counterintuitive regarding the rising spontaneous activity and excitation profiles of the current-clamp recordings, however, the development of the mean negative amplitude of synaptic input was well explained by the mean positive amplitude of synaptic events, reflecting inhibitory inputs, which was found to continuously and significantly rise from 12,43 ± 5,53 pA at the initial culturing period to 179,5 ± 26,13 pA at week 3 (figure 6E), counteracting the excitatory input and therefore decreasing the mean negative current amplitude. Interestingly, we observed a drop of the mean positive amplitude to 91,76 ± 24,08 pA in week 5 of culturing. As the amplitudes only illustrated the single most positive and negative peak points of events, we additionally again calculated the integral for each event and accounted for proportions below the baseline with negative values and proportions above the baseline with positive values to mirror inhibitory and excitatory components in the final value. Strikingly, while synaptic input was virtually non-existent in the initial culturing period, positive and negative synaptic input seemed to mostly balance each other with slightly bigger values for excitatory synaptic input until week 3, reflected by small negative values. We observed a sharp increase of the excitatory component from -9,61 ± 7,18 pAs in week 1 to -1569 ± 747 pAs in week 4, followed by a distinct drop to -234 ± 106,7 pAs in week 5 (figure 6F). The increase of synaptic input in week 4 and decrease in week 5 was in accordance with the observed development of the event length, which substantially influences the integral value. When we took the distribution of the single values into account, we could again identify clusters of small values as well as several cells displaying huge integral values due to excessive excitatory activity. All individual values are given in supplementary table 1.

**Figure 6.**
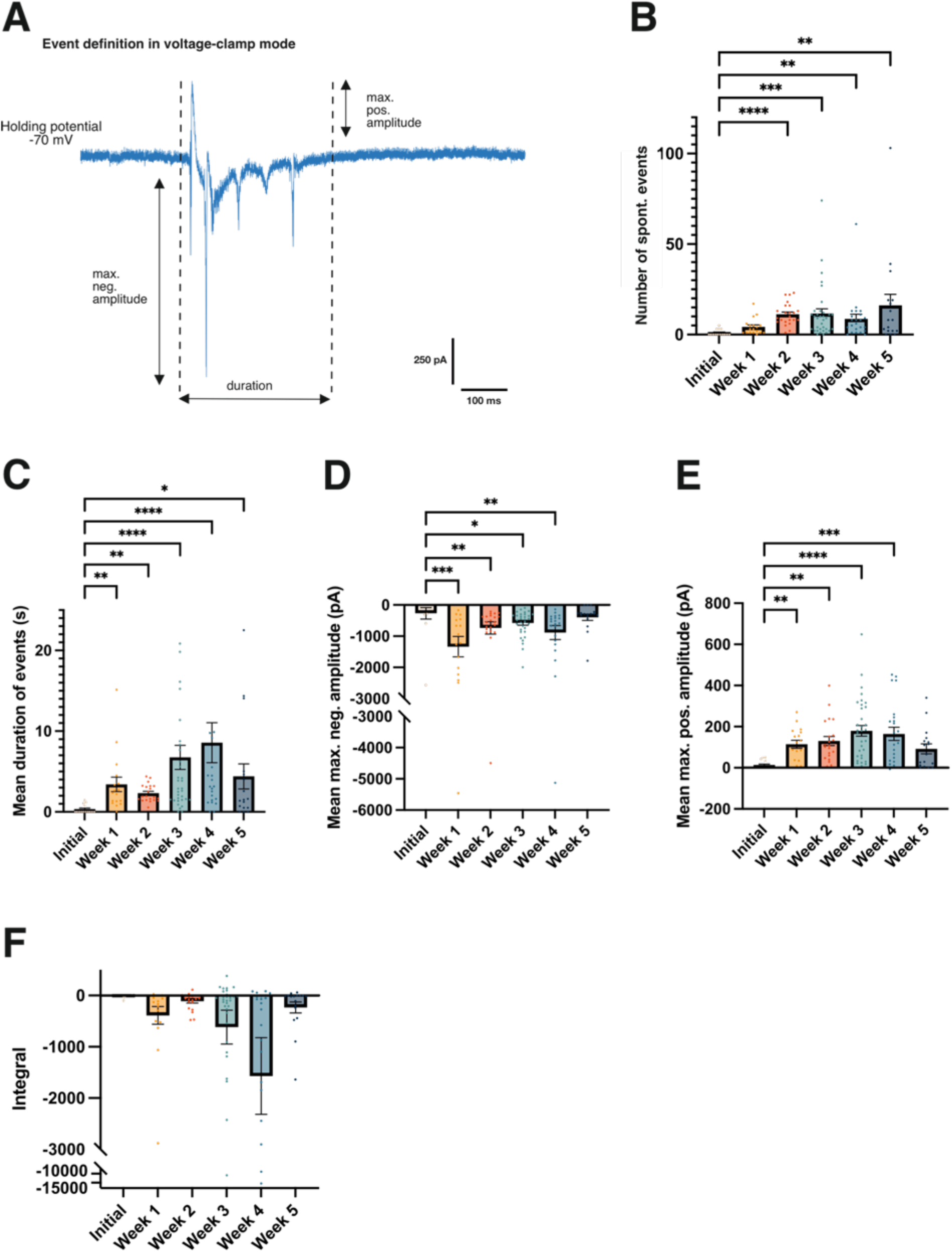
Spontaneous activity as recorded in voltage-clamp mode. (A) Definition of events during voltage-clamp recordings, event starting at reaching a threshold of 200 pA and ending when returned to baseline for at least 500 ms. Events were recorded at a holding potential of -70 mV, negative and positive amplitudes were calculated from the baseline. (B) Number of spontaneous events per cell as recorded over a recording time of three minutes. (C) Mean duration of spontaneous events. (D) Mean maximum negative amplitude. (E) Mean maximum positive amplitude. (F) Mean integral calculated for every time period, calculating area under the baseline as negative values and areas above baseline as positive values to account for excitatory/inhibitory proportions. Significance levels for all graphs: **** = p<0.0001, *** = p<0.001, ** = p<0.01, * = p<0.05 for Kruskal-Wallis-Test.

In conclusion, the development of synaptic input was coherent with the findings made in current-clamp recordings. Events gradually increased in frequency, length and complexity over the culturing period, mirroring the development of spontaneous neuronal activity. The rise of inhibitory input was observed with a slight delay to excitatory input and values became more heterogeneous and clustered into low and high activity profiles during culturing periods week 4 and 5.

### Morphological development of neocortical pyramidal neurons in culture

Next, we investigated the cellular morphology and its development over the culturing period in cells that were filled with biocytin during Patch-Clamp recordings. Cells were stained with Streptavidin, imaged on a confocal microscope and reconstructed. Per time period, we included five cells into the analysis (supplement figure 2).

All cells exhibited the typical morphological form of pyramidal cells (figure 7A) and were analyzed for their total dendritic length, number of branch points and number of terminal end points. The total dendritic length of neurons increased significantly from 1511 ± 269,6 μm at the initial culturing phase up to 5030 ± 143,2 μm in week 4, but significantly dropped to 3310 ± 286,4 μm between week 4 and week 5 (figure 7B). A similar development was observed for the number of total branch points, increasing from 24,4 ± 2,4 at initial culturing to 55 ± 3,90 by week 2, but dropping again to 37,8 ± 3,53 in week 5 (figure 7C) and number of terminal end points, which significantly increased after the initial culturing phase from 34,2 ± 3,71 to 67,4 ± 4,01 by week 2, but decreased again to 49,0 ± 3,65 in week 5 (figure 7D). A coherent development was observed when apical and basal dendrites were analyzed separately (supplement figure 3). All individual values are given in the supplement, table 1.

**Figure 7.**
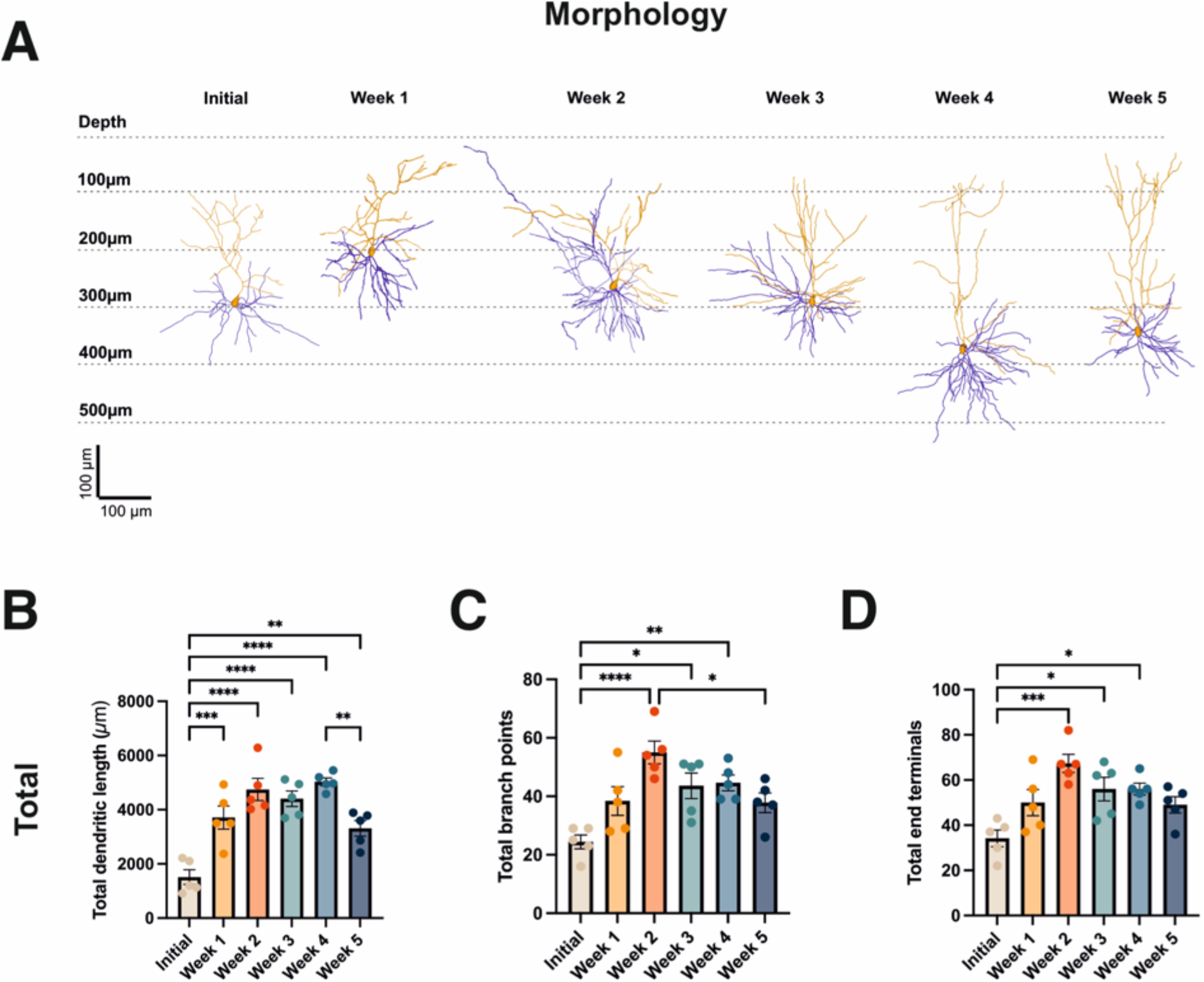
Development of pyramidal cell morphology. (A) Representative reconstructions of neurons for each culturing period, depicted at their corresponding depth as measured from the cortical surface. Apical dendrite is color-coded in orange, basal dendrites in blue. (B) Graph depicting development of total dendritic length over the culturing periods. (C) Graph depicting development of the number of total branch points over the culturing periods. (D) Graph depicting development of the number of total dendritic end terminals over the culturing periods. Significance levels for all graphs: * = p<0.05, ** = p<0.01, *** = p<0.001, **** = p<0.0001 for One-Way ANOVA test.

In conclusion, these findings mirror the ongoing maturation of neurons after the initial culturing phase up to week 2 of culturing and stability thereafter, but also again highlight the occurring changes in week 5 of culturing.

Network dynamics of neocortical organotypic brain slice cultures show transition to high-frequency oscillatory phenotype Considering our findings on the single-cell level, we were wondering how the development of spontaneous activity would reflect on a network level. Therefore, to get more insights into the overall network activity of our cortical cultured slices over time *in vitro*, we performed multi-electrode array (MEA) recordings for every time period, namely n=17(9) for initial, n=17(9) for week 1, n=13(7) for week 2, n=15(8) for week 3, n=16(8) for week 4 and n=14(7) for week 5, n = number of recordings with number of slices indicated in brackets. Recordings were performed on a 16×16 (256) electrode grid and the slice was positioned in a way that would cover the somatosensory cortical area. Afterwards, recordings were analyzed using custom written Matlab scripts (see Methods for Details) detecting every electrode that reached the individual threshold for positive or negative components of local field potentials (LFPs) and calculating their corresponding amplitudes (figure 8A, upper row).

**Figure 8.**
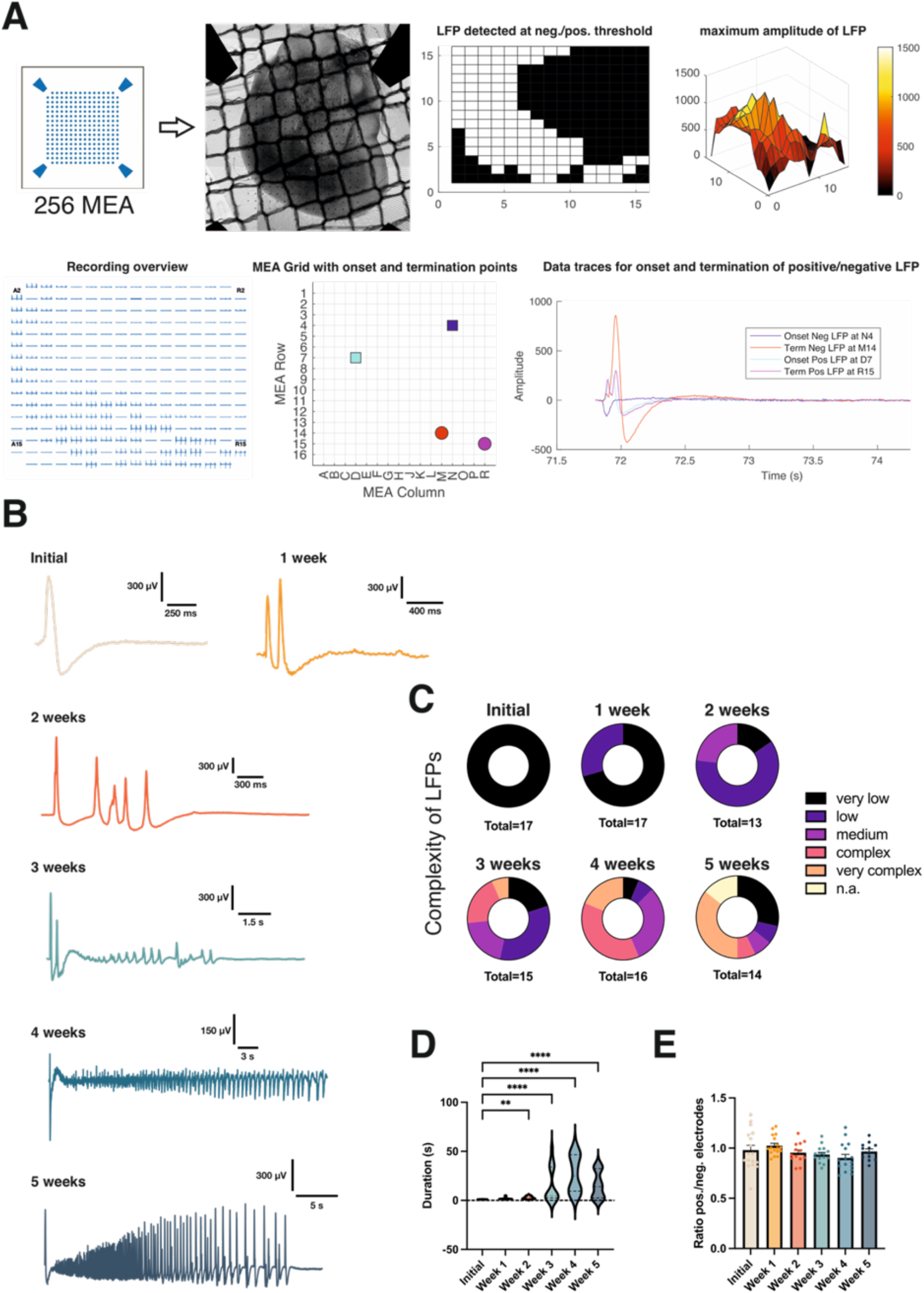
Multi-electrode array recordings of organotypic cultures. (A) Schematic representation of MEA recording and analysis pipeline. Slices were positioned on 16×16 (256) electrode grid to cover the somatosensory cortical area. Every slice was recorded for 2x five minutes and from each recording, three LFPs were analyzed at most. During analysis, electrodes reaching threshold for either positive or negative component of LFP were detected and maximum amplitude was calculated. The specific LFP was analyzed from one reference channel, representative for activity seen in the remaining electrodes as illustrated in the overview of the recording. Onset and determination points of positive and negative components were detected and waveform of the individual LFP generated. (B) Exemplary waveforms of LFPs over a time course of five weeks. (C) Development of LFP complexity over five weeks with LFPs categorized into very low (single spike), low (2-3 spikes), medium (4-5 spikes), complex (1-3 initial full sized spikes followed by low amplitude oscillation) and very complex (initial spike followed by full-sized amplitude oscillation). (D) Mean duration of LFPs for every recording over five weeks, ** = p<0.01 and **** = p<0.0001 for Kruskal-Wallis-Test. (E) Ratio of electrodes reaching positive vs. negative threshold for every recording obtained.

maximum of three LFPs per recording. Our analysis scripts detected onset and termination points for each LFP and calculated their individual amplitudes, timeframes and waveforms (figure 8A, lower row). Subsequently we analyzed which waveforms of LFPs were the most dominating throughout the slices at different time-points. We found mostly simple waveforms for the initial culturing phase, consisting of merely one positive and one negative component, followed by maturing of the activity pattern into more elaborate waveforms encompassing a higher number of different components throughout culturing week 1, week 2 and week 3, but shifting into multiple seconds long, oscillating events in week 4 and week 5 (figure 8B). Though LFP waveforms were more of a spectrum rather than clearly distinguishable subtypes, it appeared like patterns of certain complexities were dominating each culturing period. Hence, to give a better overview on which type of LFP is dominating which culturing period, we classified LFP waveform complexity by spike number and pattern, with one spike defined as consisting of both positive and negative component. LFPs were classified into complexities very low (single spike), low (2-3 spikes), medium (>3 spikes), complex (initial spike(s) followed by low amplitude oscillation, not reaching amplitude of initial spike) and very complex (initial spike(s) followed by high amplitude oscillation, reaching full amplitude of initial spike). While in the initial culturing phase, exclusively LFPs of very low complexity were observed, LFP waveforms seemed to follow a similar maturation course as observed in the other experiments. LFPs became more elaborate up until week 3 of culturing, however, complex (37.5%) and very complex (31.25%) oscillating events were taking over and dominated over half of the recordings performed by week 4 (figure 8C). Interestingly, while events of very low complexity only made up small proportions from week 2 to week 4, they suddenly reappeared as one of the dominating components in week 5 (28.57%) along with very complex oscillating events (35.71%), again illustrating the increasing heterogeneity of the cultures and their clustering into groups of low/medium basal activity and high activity, especially during week 5. This was also reflected in the duration of events, which significantly increased from 1,02 ± 0,04 s at the initial culturing phase to 15,59 ± 4,55 s in week 3 and 26,96 ± 4,75 s in week 4, but also showed distinct clustering into shorter and longer durations from week 3 to week 5 (figure 8D), resulting in a decreased mean duration of 15,72 ± 3,63 s in week 5 because of a cluster of slices exhibiting simple, short waveform LFPs as seen before in figure 8C.

Comparison of the ratio between electrodes reaching threshold for either positive or negative components of the LFP remained stable with no significant differences (figure 8E), indicating there were always both elements to every LFP, however, when we analyzed the quantitative data of LFP development, we found clear changes in LFP characteristics over time. For the positive component of the LFPs, it progressively covered significantly larger proportions of the electrode grid from 858,6 ± 48,77 μm^2^ at the initial phase to 1179 ± 48,65 μm^2^ in week 2 (figure 9A). Also, the average positive LFP amplitude increased from 0,27 ± 0,45 mV to 1,1 ± 0,14 mV by week 2 (figure 9B). The summation of positive LFPs over the grid displayed a similar development with values increasing from 20,71 ± 3,97 mV to 105,6 ± 14,89 mV by week 2 (figure 9C). A very similar trend was observed for the negative LFP component, also covering significantly more area over time in culture with values increasing from 896,1 ± 57,51 μm^2^ to 1236 ± 28,51 μm^2^ by week 2 (figure 9D). The average negative LFP amplitude increased from - 0,22 ± 0,04 mV to -0,53 ± 0,06 mV by week 4 (figure 9E) and the sum of negative LFPs from - 17,56 ± 3,52 mV to -51,07 ± 5,80 mV by week 4 (figure 9F). Interestingly, changes in average and summarized negative LFP were only significant between the initial culturing phase and week 4, which indicates that similar to the excitatory components in single-cell recordings, negative LFP components seemed to be present from the start of the culture with relative stability and only suddenly increased in late culturing weeks 4 and 5, while positive components gradually increased over the whole culturing period. All individual values are given in supplementary table 2.

**Figure 9.**
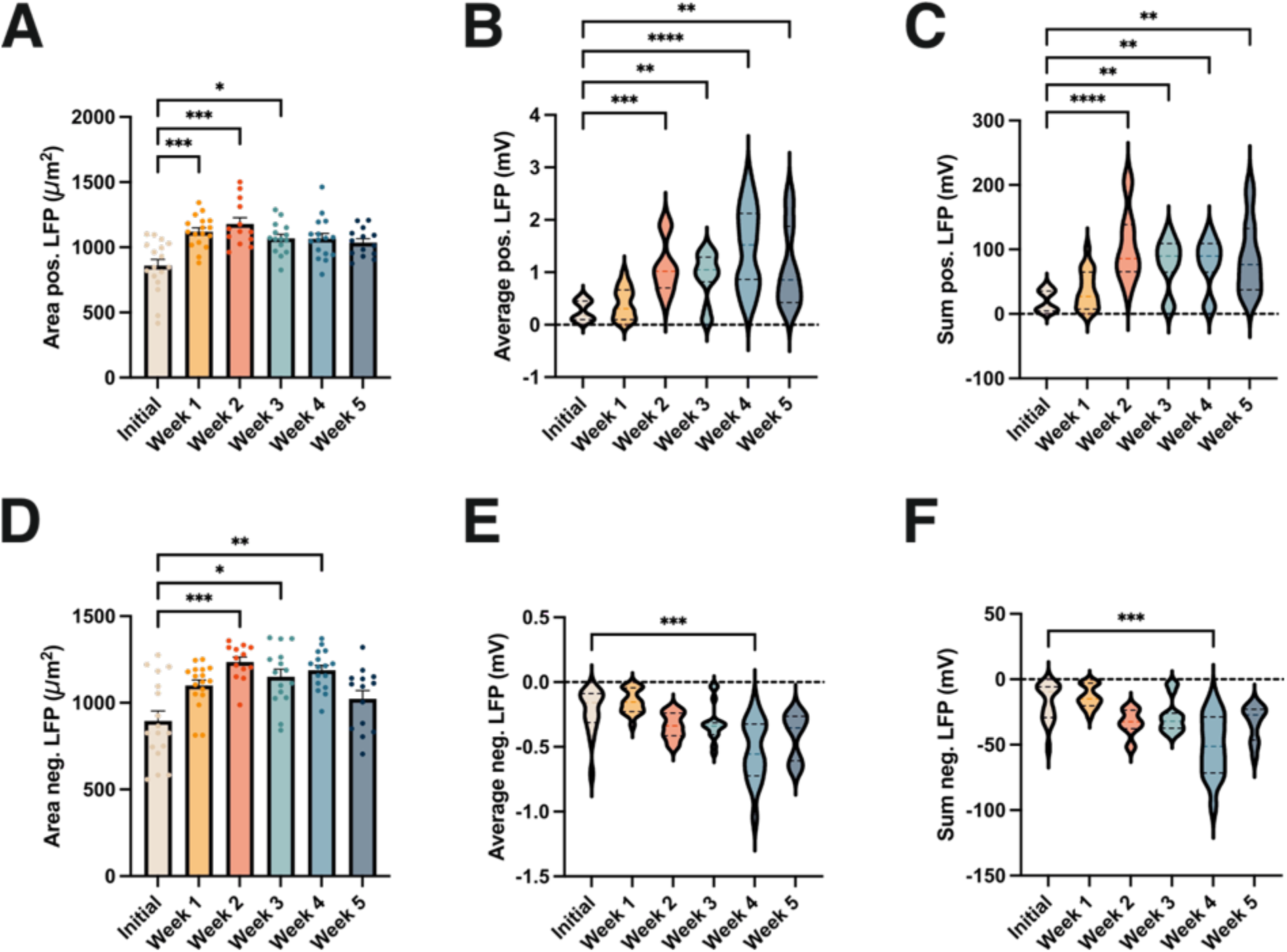
Evaluation of multi-electrode array recordings of organotypic cultures. Graphs depicting values for positive component of LFP, namely (A) area of the array covered, (B) average positive LFP and (C) sum of positive LFPs. Graphs depicting value for negative component of LFP, namely (D) area of the array covered, (E) average positive LFP and (F) sum of positive LFPs. Significance levels for all graphs: **** = p<0.0001, *** = p<0.001, ** = p<0.01 for Kruskal-Wallis-Test.

Summarized, the MEA data confirmed the maturation of spontaneous network activity into more elaborate patterns from week 1 to week 3 of organotypic neocortical cultures, but also the increase of heterogeneity and variability accompanied by the appearance of high-frequency oscillations in week 4 and 5.

### Development of the expression of *Syp, Pvalb, Hcn1, Scn1a, Scn2a*, and *Scn8a* in neocortical organotypic cultures

To gain more insight on the cellular development of our cortical oBSCs, we extracted RNA from whole hemispheres cultured as organotypic brain slices. We examined expression levels of synaptophysin as general neuronal marker, as well as parvalbumin as marker for fast-spiking interneurons. Additionally, we analyzed expression levels of *Hcn1, Scn1a, Scn2a* and *Scn8a* to investigate the development of ion conductance. Tissue was collected at 2 DIV (n=7), 7 DIV (n=7), 14 DIV (n=6), 21 DIV (n=8) and 28 DIV (n=6) respectively. Quantitative RT-PCR revealed that synaptophysin expression levels were slightly increased after the initial culturing period up until week 2 and showed a decline between week 2 and week 4 (figure 10A), indicating first signs of neuronal loss, though not significant. Parvalbumin expression increased steeply between the initial culturing period and week 3, but also decreased towards later culturing periods (figure 10B). *Hcn1* showed a trend of increasing expression between the initial culturing and week 3, decreasing again at later time points (figure 10C). *Scn1a* and *Scn8a* expression increased significantly between initial culturing and week 3, but dropped afterwards (figure 10D and F).

**Figure 10.**
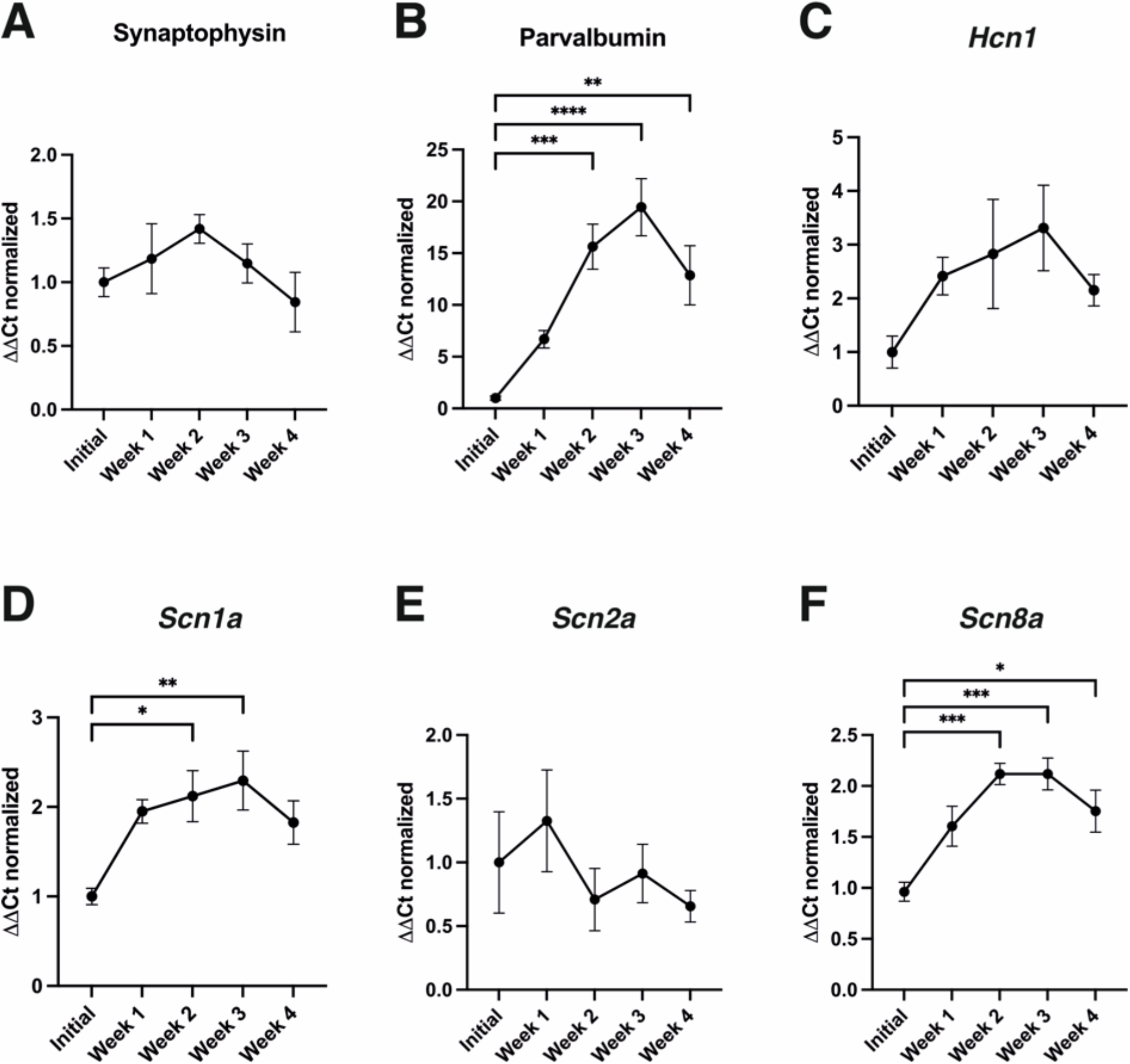
RT-PCR of markers Synaptophysin, Parvalbumin, *Hcn1, Scn1a, Scn2a, Scn8a*. Graphs depicting values for RT-PCR to evaluate development of RNA expression, namely (A) Synaptophysin, (B) Parvalbumin, (C) *Hcn1*, (D) *Scn1a*, (E) *Scn2a* and (F) *Scn8a*, all values for initial culturing period normalized to 1, expression levels of following periods all normalized in relation to values of initial phase. Significance levels for all graphs: **** = p<0.0001, *** = p<0.001, ** = p<0.01, * = p<0.05 for One-way ANOVA.

The expression curve for *Scn2a* showed high variability and was inconclusive, however, it appeared like *Scn2a* expression peaked in week 1 of culturing, but subsequently decreased and stabilized at lower levels during week 2 to week 4 (figure 10E). Especially Parvalbumin showed a clear elevation in expression between the initial culturing phase and week 3, with a near 20-fold increase in expression levels. We therefore decided to perform histological stainings on tissue sections of acute samples and cortex cultured for 7, 14 and 21 days respectively and label for NeuN and Parvalbumin.

While the general cellular and neuronal content of slices largely persisted as demonstrated by DAPI and NeuN labeling, we observed a clear upregulation of Parvalbumin-positive cells during week 1 and week 2 (figure 11A). This development was followed by a decline of expression as seen in RT-PCR, but interestingly, already seemed to appear during week 3 (figure 11B). To ensure specificity of the staining, we obtained confocal images of the stained tissue section (figure 11C). All individual values are given in supplementary table 3.

**Figure 11.**
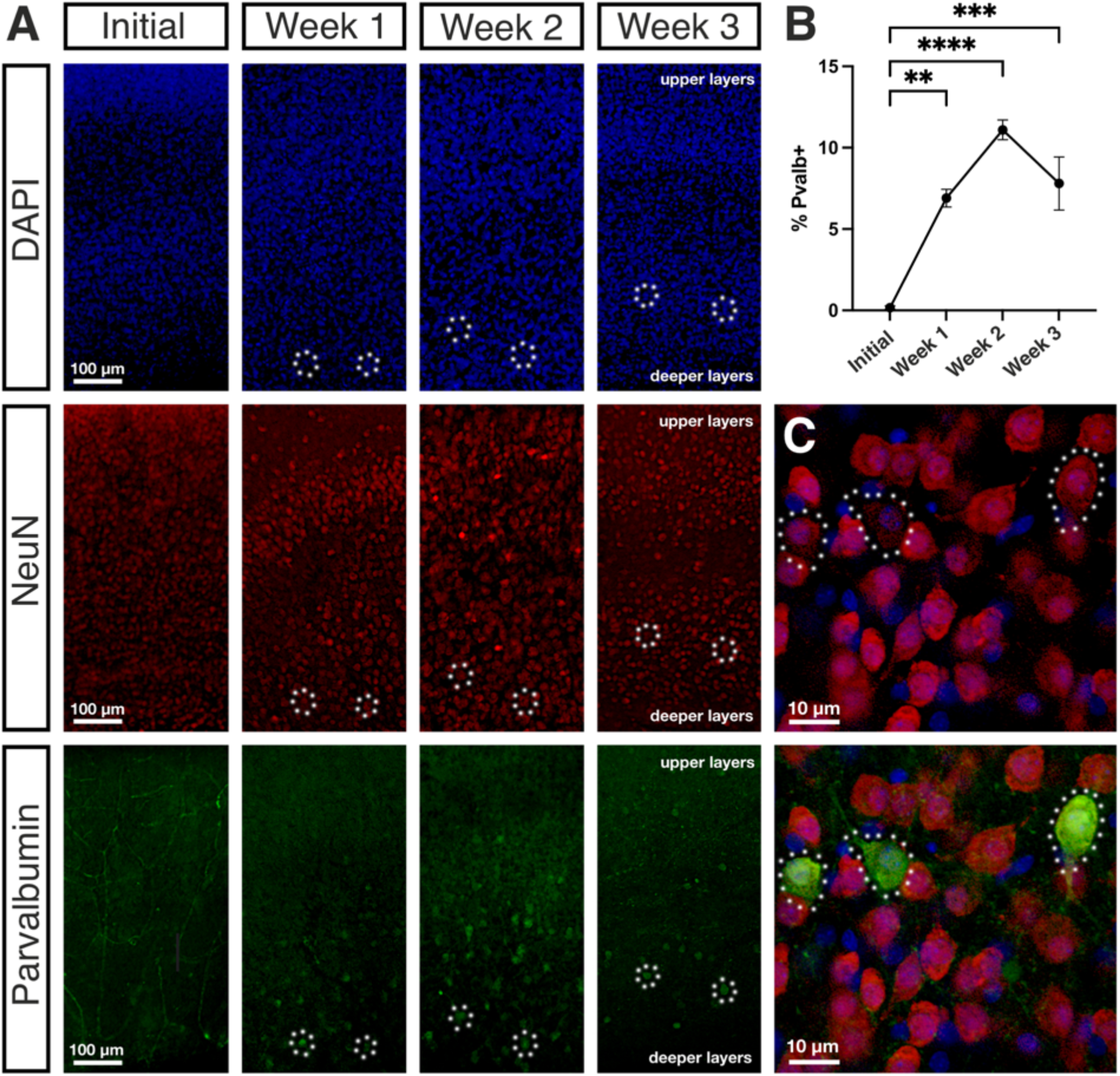
Development of parvalbumin-positive fast-spiking interneurons in organotypic cultures. (A) Stainings of cultures fixed at initial, week 1, week 2 and week 3 time periods, namely DAPI, NeuN and Parvalbumin. (B) Graphs depicting development of parvalbumin-positive neuronal population over time, ** = p<0.01, *** = p<0.001, **** = p<0.0001 for One-Way ANOVA test. (C) Specificity of Parvalbumin staining as shown by confocal images of parvalbumin-positive neurons, which were simultaneously positive for NeuN.

In conclusion, RT-PCR and staining results indicated a clear upregulation of the markers Parvalbumin, *Hcn1*, *Scn1a* and *Scn8a* between the initial culturing phase and week 3. Synaptophysin expression remained mostly stable, while *Scn2a* expression peaked in week 2 and declined afterwards. Nevertheless, nearly all markers dropped in expression by week 4, potentially indicating a general neuronal loss in late culturing phases week 4 and week 5.

## Discussion

The development of the early postnatal mouse neocortex *in vitro* concerning electrophysiological properties, morphology and RNA expression levels holds important implications for planning of experiments involving mouse cortical oBSCs. We observed that when generating these cultures from postnatal P5-6 mice, neuronal and network properties undergo significant changes over five weeks in culture. During the initial culturing phase, neurons displayed immature firing behavior characterized by a depolarization block and incomplete trains of action potentials. In the developing cortex, an increased input resistance along with a slow membrane time constant allows for the summation of excitatory postsynaptic potentials despite low network firing frequencies during early developmental stages of the cortex^45^. During the first week in culture, neurons matured and intrinsic properties as well as firing behavior became increasingly physiological as demonstrated by significantly decreasing resting membrane potential, voltage sag, input resistance and AP half-width, closely mimicking the development *in vivo*^46,47^. This maturation was also previously described in hippocampal cultures, accompanied by an increasing expression of synaptophysin^37^. Intrinsic and firing parameters were probably hugely influenced by the ion conductance dynamics of the neurons, as RNA expression levels showed a continuous increase of *Hcn1, Scn1a* and *Scn8a* expression levels between the initial culturing phase and week 3, which is also observed *in vivo*^48–50^. This maturation of ion conductance was furthermore reflected by the development of spontaneous activity, which was very rare on single-cell level in the initial culturing phase and increased over time in culture. Neurons gradually shifted from simple single spikes and short bursts to complex patterns of activity, containing elaborate up-states. This development of spontaneous activity in cultures of the somatosensory cortex along with the significant variability was described previously^39^. It is also coherent with the rise in *Hcn1* expression levels, as Hcn channels have been found to influence bursting behavior^51^. We found that spontaneous synaptic activity, while still non-existent in the initial culturing period, developed over the first 2-3 weeks in cultures with inhibitory components apparently rising with some delay to excitatory inputs. While excitation and inhibition seemed to balance each other well during the first weeks in culture, excitatory synaptic input values became more dominant during week 4 in culture, however, this likely seemed to be caused by few cells displaying heightened activity and thus, manipulating the mean value, resulting in no significance in statistical tests albeit the obvious differences. These results were coherent with the phenomenon found in current-clamp events, where cells clustered into groups of medium or low activity and high activity, again highlighting the heterogeneity of organotypic neocortical cultures in culturing week 4 and week 5. The variability of spontaneous activity on single-cell level could be attributed to the fact that network activity has been found to be initiated in L5^52^ with information in L2/3 being more sparse, variable and informative^53^. Network activity was present in all culturing phases, however, developed from a simple and short LFP waveform to high-frequency oscillations lasting multiple second by week 4 and 5. It is questionable if these events can be categorized as pathological and epileptogenic, or if they are physiological oscillations in the high-frequency range. Considering the steep increase of Parvalbumin expression levels by 20-fold as compared between the initial culturing phase and week 3, equivalent to the development seen in hippocampal cultures^54^ and *in vivo*^55^, and the retained regularity of these events (supplement figure 4), it is likely that they were not caused by runaway excitation. The oscillations observed during MEA recordings had a controlled and rhythmic character, different from the aberrant activity found in few single-cell recordings or the epileptiform phenotype described after four weeks in hippocampal cultures^11^. Up-states and network oscillations both rely on a delicate balance between excitation and inhibition, which only developed over time in culture, raising the possibility that the observed oscillations could be physiological. It has been shown that parvalbumin-positive neurons are crucial for the initiation of gamma oscillations^56^ and phase-lock to different oscillations in the hippocampus^57^, while their loss leads to a decrease in overall oscillation and especially gamma rhythm^58^. As high-frequency network events were terminated in a controlled manner, this indicates balanced excitatory and inhibitory forces. The maturation of single-cell and network properties happened independent from sensory input and therefore had to be driven intrinsically. This was also observed in sensory deprived animals, where maturation onset happened with a delay of several days, but with the same time course^59^. External electrical stimulation influences the frequency and distribution of spontaneous activity, as well as the proportion of up-states^26^. Hence, it is likely that the deprivation of external input in oBSCs is balanced by an increase in spontaneous network activity and overall excitability as observed in visually deprived animals^60^, which would explain the overall increase in excitability in culturing weeks 4 and 5 despite the network reaching a mature state by week 2. This alteration of network properties to promote excitability as a response to the loss of external input is a form of homeostatic plasticity, needed to maintain network stability and variability and depends on intact E/I balancing mechanisms^61^. Nevertheless, aberrant heightened activity and runaway excitation was observed in intracellular recordings, however, only in very few cells. The clustering into cellular and network phenotypes with either very basal or very high activity, especially observed in week 5 of culturing, could be due to a general heterogeneity in slice viability or potentially even differential changes in transcriptomic profiles influencing homeostatic plasticity and activity, such as the transcription factor family PARbZip^62^. A certain degree of neuronal loss definitely takes place during weeks 4 and 5, evidenced by the collective drop of RNA expression levels in these culturing periods. Moreover, an indication of potential degradation in week 5 was also found in morphological analysis by the significant loss of neuronal complexity. However, the general dendritic arborization of pyramidal neurons in culture closely mimics the *in vivo* maturation process during the first postnatal week^63^ and both *in vitro* and *in vivo* results indicate that the final dendritic configuration is established after this time frame. This is highlighted by the fact that coherent morphological changes could not be found in studies investigating mice from P10 onward^46^.

## Conclusion

In this study, we provided a holistic characterization of the development of the early postnatal mouse neocortex over five weeks *in vitro*. In summary, the immature cortex underwent a maturation similar as observed *in vivo* in terms of intrinsic and firing properties, network activity and RNA expression levels. Cortical oBSCs therefore approach a mature state after approximately two weeks *in vitro*. In conclusion, experiments aiming to investigate developmental phenomena of the cortex are best performed from 0-14 days *in vitro*. In contrast, investigations in need of an adult configuration of the network, e.g. simulations of diseases of the adult brain, will find the cortical network to be most stable and physiologically mature in weeks 2 and 3 of culturing. Beyond this time frame, rising variability, heterogeneity, excitability and potential neuronal loss have to be considered, however, cortical cultures aged 4 or 5 weeks could potentially serve as models for network oscillations.

## Acknowledgements

This work was funded by the Chan Zuckerberg Initiative Collaborative Pairs Pilot Project Awards (Phase 1 & Phase 2) and by the German Research Foundation (DFG/FNR INTER research unit FOR2715).

This work was supported by the "Confocal Microscopy Facility", a core facility of the Interdisciplinary Center for Clinical Research (IZKF) Aachen within the Faculty of Medicine at RWTH Aachen University.

## Notes

### Competing Interest Statement

The authors have declared no competing interest.

