## Supplement data for "Temporal Dynamics of Neocortical Development in Organotypic Mouse Cultures: A Comprehensive Analysis"

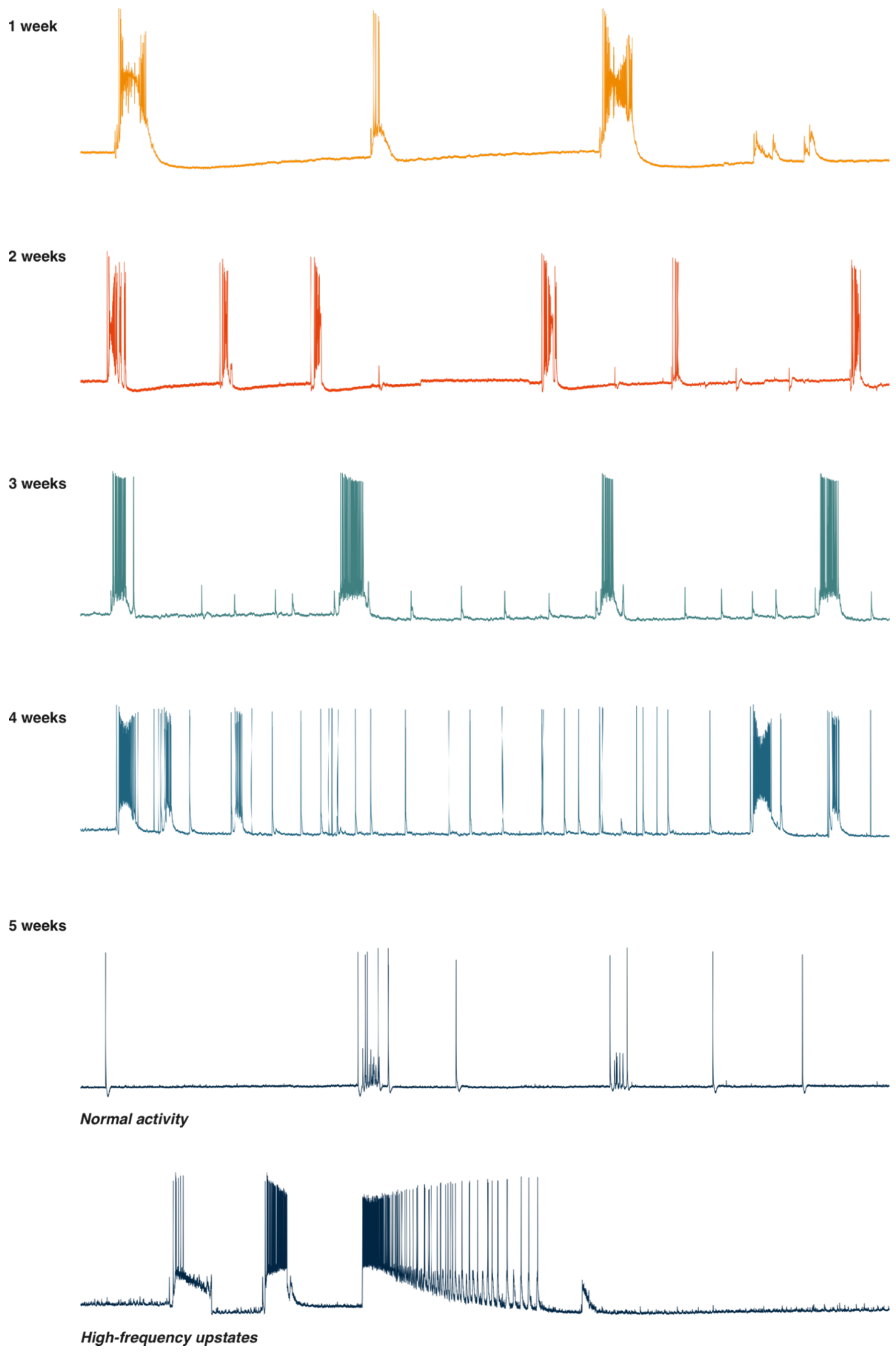


Supplement, Figure 1. **Development of spontaneous activity in whole-cell Patch-Clamp recordings.** Representative continuous recordings of three minutes depicted for every culturing period. Up-states were found from one week in culture onward and cells generally depicted diverse patterns of activity, containing single spikes, bursts and up-states. Week 5 neurons clustered in low/medium basal activity phenotypes and neurons exhibiting high-frequency up-states.

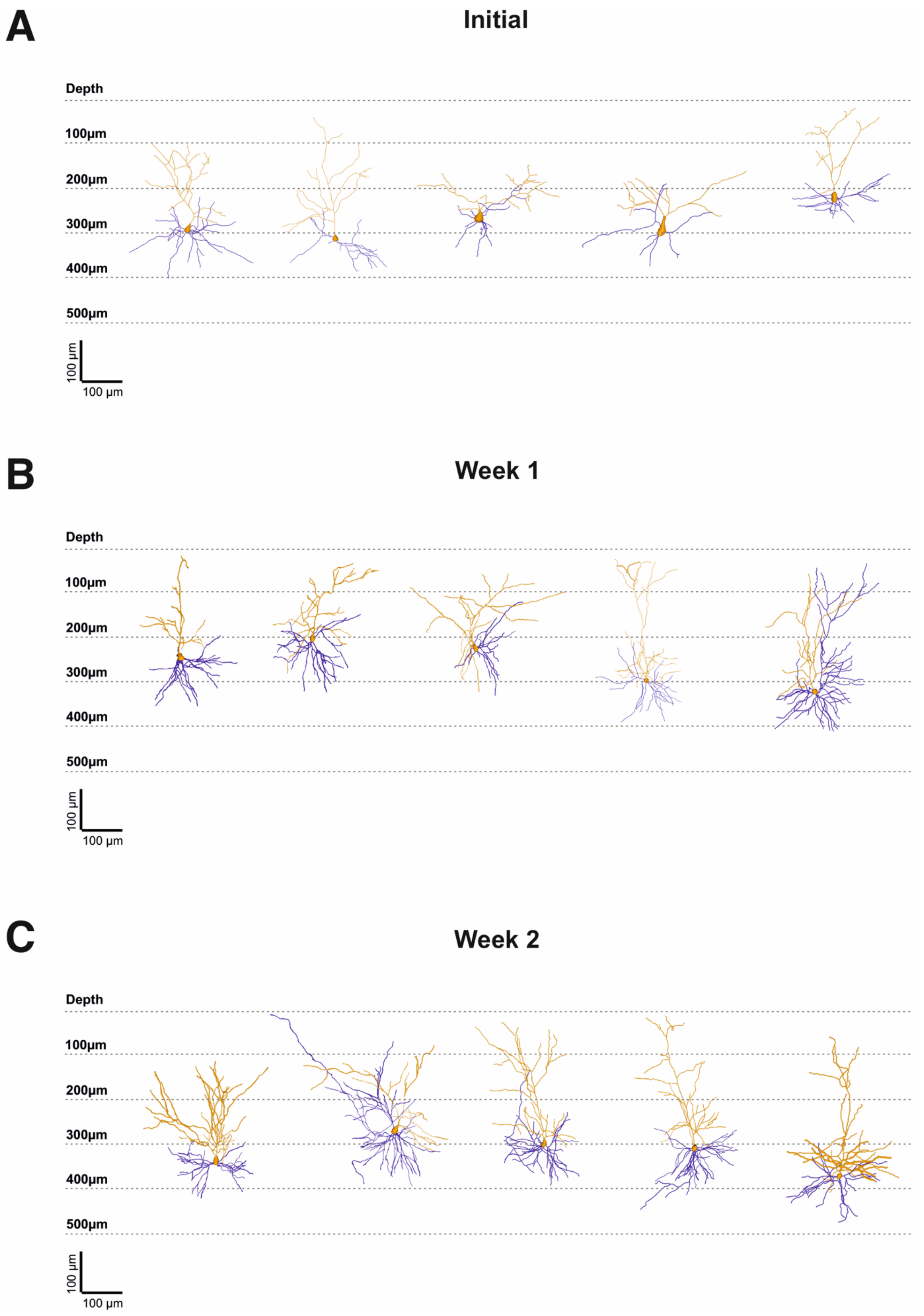
A
Initial
Depth
100µm
200µm
300µm
400µm
500µm
100 µm
100 µm
B
Week 1
Depth
100µm
200µm
300µm
400µm
500µm
100 µm
100 µm
C
Week 2
Depth
100µm
200µm
300µm
400µm
500µm
100 µm
100 µm

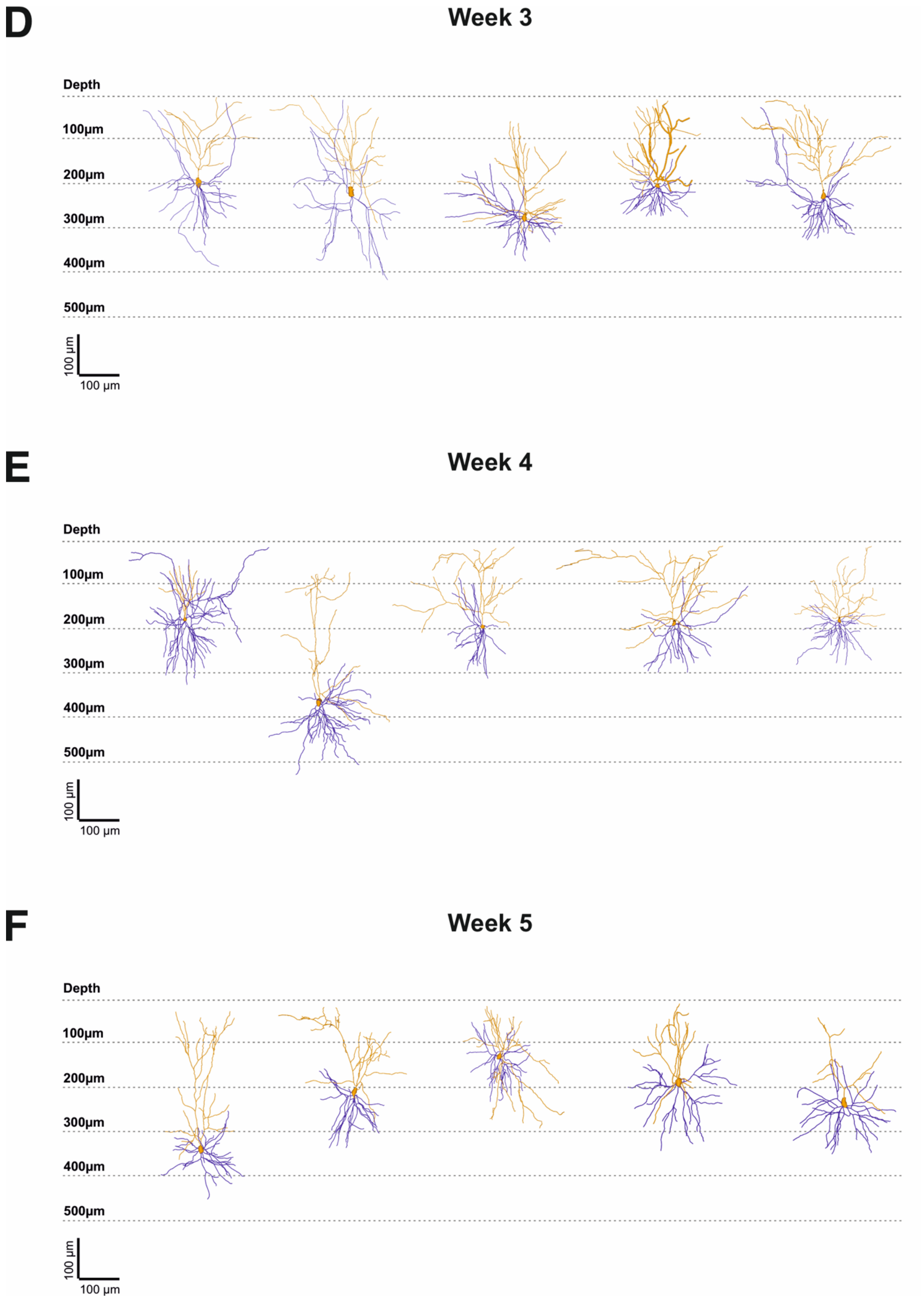


Supplement, Figure 2. **Pyramidal neurons included in morphological analysis.** Five neurons per culturing period were included in morphological analysis, depicted with corresponding depth from cortical surface. Apical dendrite is color-coded in orange, basal dendrites in blue.

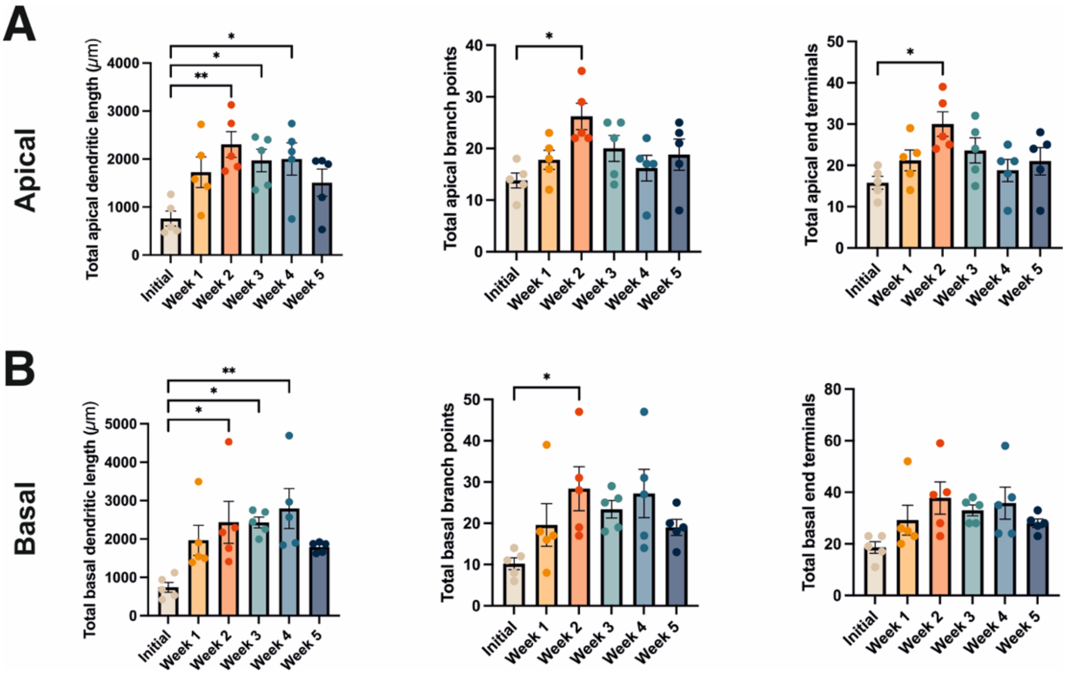


Supplement, Figure 3. **Development of morphological properties of apical and basal dendrites.** Analysis of the development of dendritic length, dendritic branch points and dendritic end terminals over five weeks. (A) Values depicted for the apical dendrite section of neurons. (B) Values depicted for the basal dendrites of neurons. Significance levels for all graphs: * = p<0.05, ** = p<0.01, *** = p<0.001, **** = p<0.0001 for One-Way ANOVA test.

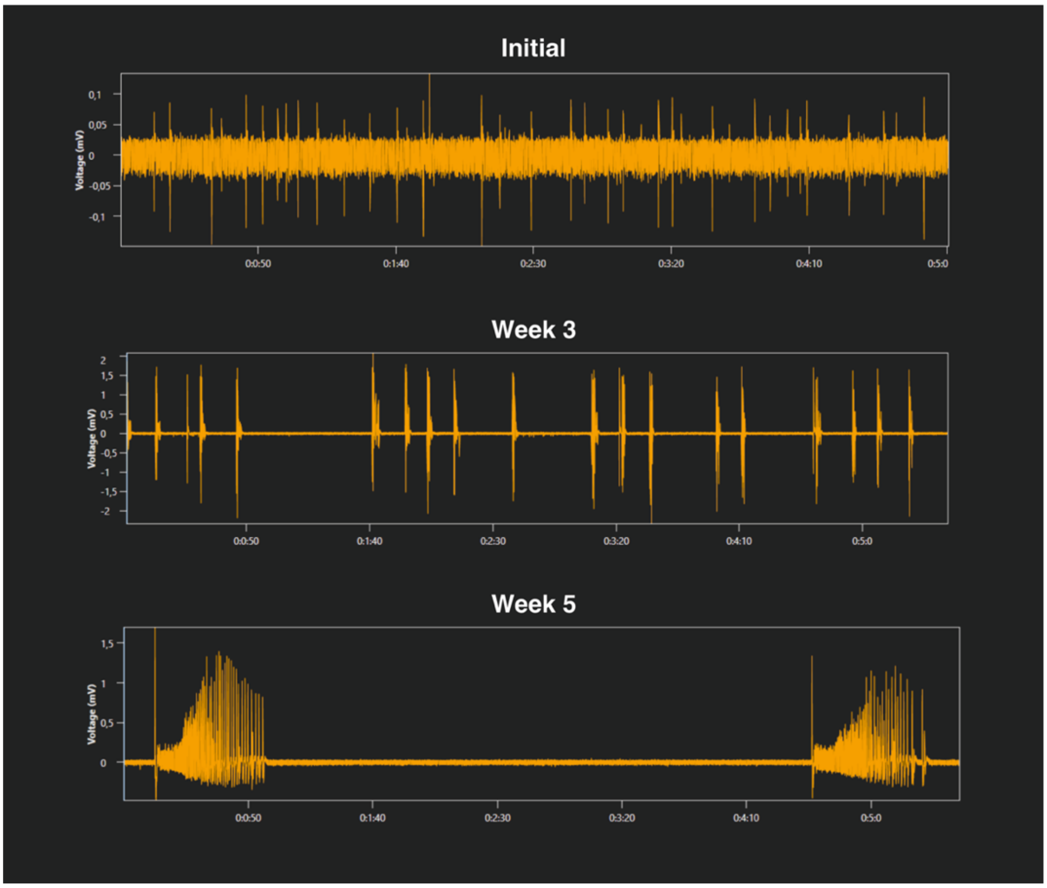


Supplement, Figure 4. **Representative traces from MEA recordings.** Representative channel of MEA recordings of five minutes length depicted for culturing periods initial, week 3 and week 5. Events gradually increase in amplitude, length and complexity, but regularity and rhythmicity was preserved.

**Passive & firing properties**

| Parameter | Unit | Initial (n=21) | Week 1 (n=28) | Week 2 (n=36) | Week 3 (n=45) | Week 4 (n=30) | Week 5 (n=19) |
|---|---|---|---|---|---|---|---|
| **Membrane potential** | mV | -73,65 ± 0,96 | -74,52 ± 1,22 | -81,19 ± 1,72 | -78,52 ± 1,42 | -81,07 ± 1,79 | -80,96 ± 2,02 |
| **Voltage sag** | mV | 9,065 ± 1,04 | 3,141 ± 0,47 | 1,724 ± 0,27 | 1,687 ± 0,31 | 1,654 ± 0,50 | 2,473 ± 0,61 |
| **Input resistance** | MΩ | 338,2 ± 29,19 | 168 ± 9,91 | 139,7 ± 7,38 | 148,3 ± 9,99 | 138,7 ± 8,82 | 146,4 ± 11,2 |
| **Comparison Current/nAPs** | AUC | 2472 ± 252,4 | 1830 ± 277,2 | 1148 ± 207,4 | 2059 ± 409,1 | 1853 ± 357,1 | 2430 ± 624,9 |
| **AP width** | ms | 2,615 ± 0,15 | 2,439 ± 0,11 | 2,253 ± 0,11 | 1,858 ± 0,10 | 1,305 ± 0,04 | 1,218 ± 0,04 |
| **Comparison Inst. Frequency** | AUC | 269,3 ± 31,04 | 159,7 ± 17,51 | 124,5 ± 16,07 | 361,8 ± 76,05 | 259,7 ± 58,85 | 528,8 ± 65,28 |

**Mean (rank) difference**

| Parameter | In. vs. W1 | In. vs. W2 | In. vs. W3 | In. vs. W4 | In. vs. W5 | W1 vs. W2 | W1 vs. W3 | W1 vs. W4 | W1 vs. W5 | W2 vs. W3 | W2 vs. W4 | W2 vs. W5 | W3 vs. W4 | W3 vs. W5 | W4 vs. W5 |
|---|---|---|---|---|---|---|---|---|---|---|---|---|---|---|---|
| Membrane potential | 0,88 (ns) | 7,54 (*) | 4,87 (ns) | 7,43 (*) | 7,32 (ns) | 6,66 (*) | 3,99 (ns) | 6,55 (ns) | 6,44 (ns) | -2,67 (ns) | -0,12 (ns) | -0,22 (ns) | 2,56 (ns) | 2,45 (ns) | -0,11 (ns) |
| Voltage sag | 5,92 (****) | 7,34 (****) | 7,63 (****) | 7,98 (****) | 6,59 (****) | 1,42 (ns) | 1,70 (*) | 2,06 (*) | 0,67 (ns) | 0,29 (ns) | 0,64 (ns) | -0,75 (ns) | 0,35 (ns) | -1,03 (ns) | -1,39 (ns) |
| Input resistance | 170,2 (****) | 198,5 (****) | 189,9 (****) | 199,5 (****) | 191,8 (****) | 28,26 (ns) | 19,68 (ns) | 29,32 (ns) | 21,63 (ns) | -8,58 (ns) | 1,06 (ns) | -6,63 (ns) | 9,64 (ns) | 1,95 (ns) | -7,69 (ns) |
| Comparison Current/nAPs | 19,3 (ns) | 38,49 (**) | 21,68 (ns) | 20,53 (ns) | 13,13 (ns) | 19,18 (ns) | 2,38 (ns) | 1,23 (ns) | -6,17 (ns) | -16,8 (ns) | -17,96 (ns) | -25,35 (ns) | -1,15 (ns) | -8,55 (ns) | -7,40 (ns) |
| AP width | 0,09 (ns) | 0,54 (****) | 0,95 (****) | 1,37 (****) | 1,40 (****) | 0,45 (***) | 0,86 (****) | 1,28 (****) | 1,31 (****) | 0,41 (***) | 0,82 (****) | 0,86 (****) | 0,41 (**) | 0,44 (**) | 0,03 (ns) |
| Comparison Inst. Frequency | 19,32 (ns) | 26,85 (ns) | 2,35 (ns) | 13,58 (ns) | -20,58 (ns) | 7,52 (ns) | -16,98 (ns) | -5,74 (ns) | -39,9 (**) | -24,5 (ns) | -13,27 (ns) | -47,43 (***) | 11,23 (ns) | -22,93 (ns) | -34,16 (*) |

**Spontaneous activity (single-cell, current-clamp)**

| Parameter | Unit | Initial (n=17) | Week 1 (n=25) | Week 2 (n=30) | Week 3 (n=36) | Week 4 (n=23) | Week 5 (n=17) |
|---|---|---|---|---|---|---|---|
| **Event duration** | s | 0,2727 ± 0,16 | 2,516 ± 0,45 | 2,371 ± 0,43 | 5,582 ± 1,33 | 4,606 ± 1,89 | 7,972 ± 4,72 |
| **Number of events** | # | 0,8824 ± 0,46 | 5,08 ± 1,62 | 8,967 ± 1,52 | 3,806 ± 1,01 | 9,043 ± 3,05 | 9,706 ± 3,83 |
| **Time in active state** | % | 0,5124 ± 0,27 | 4,684 ± 0,87 | 11,81 ± 2,48 | 8,38 ± 1,55 | 9,746 ± 1,90 | 13,04 ± 4,92 |
| **APs per cell** | # | 1,235 ± 0,63 | 28,8 ± 7,23 | 63,7 ± 18,58 | 53,58 ± 17,12 | 56,78 ± 22,87 | 108,2 ± 64,22 |
| **APs/event** | # | 0,3529 ± 0,18 | 7,244 ± 1,98 | 4,88 ± 1,10 | 20,2 ± 12,51 | 16,15 ± 13,33 | 25,14 ± 20,41 |
| **Integral** | mVs | 1,232 ± 0,93 | 34,97 ± 10,98 | 17,29 ± 6,09 | 66,19 ± 20,38 | 79,85 ± 50,43 | 95,11 ± 75,48 |

**Mean (rank) difference**

| Parameter | In. vs. W1 | In. vs. W2 | In. vs. W3 | In. vs. W4 | In. vs. W5 | W1 vs. W2 | W1 vs. W3 | W1 vs. W4 | W1 vs. W5 | W2 vs. W3 | W2 vs. W4 | W2 vs. W5 | W3 vs. W4 | W3 vs. W5 | W4 vs. W5 |
|---|---|---|---|---|---|---|---|---|---|---|---|---|---|---|---|
| Event duration | -49,88 (**) | -48,04 (**) | -52,81 (***) | -50,91 (**) | -44,74 (*) | 1,84 (ns) | -2,93 (ns) | -1,03 (ns) | 5,14 (ns) | -4,77 (ns) | -2,86 (ns) | 3,31 (ns) | 1,90 (ns) | 8,07 (ns) | 6,17 (ns) |
| Number of events | -40,66 (*) | -60,99 (**** | -30,98 (ns) | -48,13 (**) | -44,21 (*) | -20,33 (ns) | 9,69 (ns) | -7,47 (ns) | -3,54 (ns) | 30,01 (ns) | 12,86 (ns) | 16,78 (ns) | -17,15 (ns) | -13,23 (ns) | 3,92 (ns) |
| Time in active state | -38,69 (ns) | -62,49 (**** | -47,73 (**) | -56,81 (***) | -52,79 (**) | -23,8 (ns) | -9,05 (ns) | -18,13 (ns) | -14,11 (ns) | 14,76 (ns) | 5,68 (ns) | 9,70 (ns) | -9,08 (ns) | -6,06 (ns) | 4,02 (ns) |
| APs per cell | -49,66 (**) | -57,82 (***) | -43,21 (**) | -47,15 (**) | -50,41 (**) | -8,16 (ns) | 6,45 (ns) | 2,51 (ns) | -0,76 (ns) | 14,61 (ns) | 10,67 (ns) | 7,41 (ns) | -3,94 (ns) | -7,20 (ns) | -3,26 (ns) |
| APs/event | -57,49 (***) | -51,43 (***) | -48,97 (**) | -42,51 (*) | -46,79 (*) | 6,06 (ns) | 8,52 (ns) | 14,98 (ns) | 10,7 (ns) | 2,46 (ns) | 8,92 (ns) | 4,64 (ns) | 6,47 (ns) | 2,18 (ns) | -4,29 (ns) |
| Integral | -52,5 (**) | -38,22 (*) | -47,04 (**) | -38,87 (ns) | -36,03 (ns) | 14,28 (ns) | 5,45 (ns) | 13,62 (ns) | 16,47 (ns) | -8,82 (ns) | -0,65 (ns) | 2,19 (ns) | 8,18 (ns) | 11,01 (ns) | 2,84 (ns) |

**Spontaneous activity (single-cell, voltage-clamp)**

| Parameter | Unit | Initial (n=14) | Week 1 (n=17) | Week 2 (n=21) | Week 3 (n=34) | Week 4 (n=23) | Week 5 (n=17) |
|---|---|---|---|---|---|---|---|
| **Number of events** | # | 1 ± 0,46 | 3,882 ± 1,03 | 11,1 ± 1,39 | 11,68 ± 2,53 | 8,696 ± 2,57 | 16,12 ± 6,15 |
| **Event duration** | s | 0,3176 ± 0,14 | 3,402 ± 0,90 | 2,315 ± 0,23 | 6,76 ± 1,49 | 8,573 ± 2,48 | 4,396 ± 1,56 |
| **Max. neg. amplitude** | pA | -267,9 ± 193,2 | -1337 ± 324,8 | -736,3 ± 195,4 | -575,3 ± 74,16 | -884,9 ± 225,9 | -394,9 ± 101,8 |
| **Max. pos. Amplitude** | pA | 12,43 ± 5,53 | 114,3 ± 19,54 | 130,3 ± 22,11 | 179,5 ± 26,13 | 164,8 ± 32,06 | 91,76 ± 24,08 |
| **Integral** | pAs | -9,612 ± 7,176 | -387,7 ± 171,3 | -111,4 ± 34,83 | -616,7 ± 329,2 | -1569 ± 747 | -234 ± 106,7 |

**Mean (rank) difference**

| Parameter | In. vs. W1 | In. vs. W2 | In. vs. W3 | In. vs. W4 | In. vs. W5 | W1 vs. W2 | W1 vs. W3 | W1 vs. W4 | W1 vs. W5 | W2 vs. W3 | W2 vs. W4 | W2 vs. W5 | W3 vs. W4 | W3 vs. W5 | W4 vs. W5 |
|---|---|---|---|---|---|---|---|---|---|---|---|---|---|---|---|
| Number of events | -25,26 (ns) | -63,23 (**** | -51,41 (***) | -43,86 (**) | -49,23 (**) | -37,97 (*) | -26,15 (ns) | -18,6 (ns) | -23,97 (ns) | 11,82 (ns) | 19,37 (ns) | 14,0 (ns) | 7,55 (ns) | 2,18 (ns) | -5,37 (ns) |
| Event duration | -47,32 (**) | -47,26 (**) | -59,32 (**** | -61,69 (**** | -41,61 (*) | 0,06 (ns) | -12,0 (ns) | -14,37 (ns) | 5,71 (ns) | -12,06 (ns) | -14,42 (ns) | 5,65 (ns) | -2,37 (ns) | 17,71 (ns) | 20,07 (ns) |
| Max. neg. amplitude | 59,14 (***) | 44,48 (**) | 38,55 (*) | 46,16 (**) | 19,02 (ns) | -14,66 (ns) | -20,59 (ns) | -12,98 (ns) | -40,12 (*) | -5,93 (ns) | 1,68 (ns) | -25,46 (ns) | 7,61 (ns) | -19,53 (ns) | -27,13 (ns) |
| Max. pos. Amplitude | -45,33 (**) | -48,86 (**) | -56,63 (**** | -50,5 (***) | -33,04 (ns) | -3,53 (ns) | -11,29 (ns) | -5,17 (ns) | 12,29 (ns) | -7,77 (ns) | -1,64 (ns) | 15,82 (ns) | 6,13 (ns) | 23,59 (ns) | 17,46 (ns) |
| Integral | 34,89 (ns) | 24,05 (ns) | 25,39 (ns) | 34,53 (ns) | 23,72 (ns) | -10,85 (ns) | -9,5 (ns) | -0,367 (ns) | -11,18 (ns) | 1,35 (ns) | 10,48 (ns) | -0,33 (ns) | 9,13 (ns) | -1,68 (ns) | -10,81 (ns) |

**Morphology**

| Parameter | Unit | Initial (n=5) | Week 1 (n=5) | Week 2 (n=5) | Week 3 (n=5) | Week 4 (n=5) | Week 5 (n=5) |
|---|---|---|---|---|---|---|---|
| **Dendritic length (total)** | µm | 1511 ± 269,6 | 3708 ± 427,2 | 4745 ± 411,4 | 4405 ± 287,4 | 5030 ± 143,2 | 3310 ± 286,4 |
| **Branch points (total)** | # | 24,4 ± 2,4 | 38,4 ± 4,93 | 55 ± 3,90 | 43,6 ± 4,38 | 44,6 ± 2,69 | 37,8 ± 3,53 |
| **End terminals (total** | # | 34,2 ± 3,71 | 50 ± 5,79 | 67,4 ± 4,01 | 56 ± 5,23 | 56 ± 2,59 | 49 ± 3,65 |
| **Dendritic length (apical)** | µm | 760,4 ± 154,9 | 1725 ± 319,1 | 2304 ± 268,9 | 1969 ± 235,9 | 2000 ± 336,4 | 1509 ± 283,3 |
| **Branch points (apical)** | # | 13,8 ± 1,46 | 17,8 ± 1,86 | 26,2 ± 2,56 | 20 ± 2,53 | 16,2 ± 2,48 | 18,8 ± 3,01 |
| **End terminals (apical)** | # | 15,8 ± 1,56 | 21,2 ± 2,52 | 30 ± 3,0 | 23,6 ± 3,04 | 18,8 ± 2,69 | 21 ± 3,33 |
| **Dendritic length (basal)** | µm | 740 ± 129,1 | 1966 ± 391,4 | 2434 ± 547,3 | 2429 ± 141,3 | 2793 ± 520 | 1789 ± 61,98 |
| **Branch points (basal)** | # | 10,2 ± 1,43 | 19,6 ± 5,16 | 28,4 ± 5,34 | 23,4 ± 2,11 | 27,2 ± 5,85 | 19 ± 1,92 |
| **End terminals (basal)** | # | 18,6 ± 2,23 | 29,2 ± 5,79 | 37,8 ± 6,21 | 33 ± 2,10 | 35,8 ± 6,23 | 28 ± 1,61 |

**Mean (rank) difference**

| Parameter | In. vs. W1 | In. vs. W2 | In. vs. W3 | In. vs. W4 | In. vs. W5 | W1 vs. W2 | W1 vs. W3 | W1 vs. W4 | W1 vs. W5 | W2 vs. W3 | W2 vs. W4 | W2 vs. W5 | W3 vs. W4 | W3 vs. W5 | W4 vs. W5 |
|---|---|---|---|---|---|---|---|---|---|---|---|---|---|---|---|
| Dendritic length (total) | -2197 (***) | -3235 (****) | -2894 (****) | -3519 (****) | -1799 (**) | -1037 (ns) | -697,2 (ns) | -1322 (ns) | 398,4 (ns) | 340,2 (ns) | -284,2 (ns) | 1436 (*) | -624,4 (ns) | 1096 (ns) | 1720 (**) |
| Branch points (total) | -14,0 (ns) | -30,6 (****) | -19,2 (*) | -20,2 (**) | -13,4 (ns) | -16,6 (*) | -5,2 (ns) | -6,2 (ns) | 0,6 (ns) | 11,4 (ns) | 10,4 (ns) | 17,2 (*) | -1,0 (ns) | 5,8 (ns) | 6,8 (ns) |
| End terminals (total | -15,8 (ns) | -33,2 (***) | -21,8 (*) | -21,8 (*) | -14,8 (ns) | -17,4 (ns) | -6,0 (ns) | -6,0 (ns) | 1,0 (ns) | 11,4 (ns) | 11,4 (ns) | 18,4 (ns) | 0 (ns) | 7,0 (ns) | 7,0 (ns) |
| Dendritic length (apical) | -965 (ns) | -1544 (**) | -1209 (*) | -1240 (*) | -749 (ns) | -579 (ns) | -244 (ns) | -274,8 (ns) | 216 (ns) | 335 (ns) | 304,2 (ns) | 795 (ns) | -30,8 (ns) | 460 (ns) | 490,8 (ns) |
| Branch points (apical) | -4 (ns) | -12,4 (*) | -6,2 (ns) | -2,4 (ns) | -5 (ns) | -8,4 (ns) | -2,2 (ns) | 1,6 (ns) | -1 (ns) | 6,2 (ns) | 10 (ns) | 7,4 (ns) | 3,8 (ns) | 1,2 (ns) | -2,6 (ns) |
| End terminals (apical) | -5,4 (ns) | -14,2 (*) | -7,8 (ns) | -3 (ns) | -5,2 (ns) | -8,8 (ns) | -2,4 (ns) | 2,4 (ns) | 0,2 (ns) | 6,4 (ns) | 11,2 (ns) | 9 (ns) | 4,8 (ns) | 2,6 (ns) | -2,2 (ns) |
| Dendritic length (basal) | -1226 (ns) | -1694 (*) | -1689 (*) | -2053 (**) | -1049 (ns) | -468 (ns) | -463,2 (ns) | -827,4 (ns) | 177,2 (ns) | 4,8 (ns) | -359,4 (ns) | 645,2 (ns) | -364,2 (ns) | 640,4 (ns) | 1005 (ns) |
| Branch points (basal) | -9,4 (ns) | -18,2 (*) | -13,2 (ns) | -17 (ns) | -8,8 (ns) | -8,8 (ns) | -3,8 (ns) | -7,6 (ns) | 0,6 (ns) | 5 (ns) | 1,2 (ns) | 9,4 (ns) | -3,8 (ns) | 4,4 (ns) | 8,2 (ns) |
| End terminals (basal) | -10,6 (ns) | -19,2 (ns) | -14,4 (ns) | -17,2 (ns) | -9,4 (ns) | -8,6 (ns) | -3,8 (ns) | -6,6 (ns) | 1,2 (ns) | 4,8 (ns) | 2 (ns) | 9,8 (ns) | -2,8 (ns) | 5 (ns) | 7,8 (ns) |

Supplement, Table 1. **Values of single-cell recordings and morphology.** All values given for every time period with corresponding unit, n-numbers and standard error of the mean. Each time period statistically compared with each other period and mean rank difference given with significance levels. Significance levels: ns = not significant, **** = p<0.0001, *** = p<0.001, ** = p<0.01, * = p<0.05

**Network activity (MEA recordings)**

| Parameter | Unit | Initial (n=17/9) | Week 1 (n=17/9) | Week 2 (n=13/7) | Week 3 (n=15/8) | Week 4 (n=16/8) | Week 5 (n=14/7) |
|---|---|---|---|---|---|---|---|
| **Event duration** | s | 1,023 ± 0,04 | 1,698 ± 0,23 | 2,909 ± 0,36 | 15,59 ± 4,55 | 26,96 ± 4,75 | 15,72 ± 3,63 |
| **Ratio pos./neg. electrodes** | - | 0,9809 ± 0,05 | 1,024 ± 0,02 | 0,9538 ± 0,03 | 0,9372 ± 0,02 | 0,9025 ± 0,04 | 0,9671 ± 0,03 |
| **Area pos. LFPs** | µm2 | 858,6 ± 48,77 | 1119 ± 29,74 | 1179 ± 48,65 | 1069 ± 31,06 | 1065 ± 40,59 | 1036 ± 28,95 |
| **Average pos. LFP** | mV | 0,27 ± 0,45 | 0,41 ± 0,07 | 1,1 ± 0,14 | 0,94 ± 0,12 | 1,47 ± 0,19 | 1,08 ± 0,20 |
| **Sum of pos. LFPs** | mV | 20,71 ± 3,97 | 38,19 ± 7,26 | 105,6 ± 14,89 | 78,82 ± 10,09 | 78,82 ± 10,09 | 87,2 ± 14,98 |
| **Area neg. LFPs** | µm2 | 896,1 ± 57,51 | 1100 ± 31,39 | 1236 ± 28,51 | 1151 ± 43,1 | 1188 ± 28,47 | 1023 ± 47,81 |
| **Average neg. LFP** | mV | -0,2231 ± 0,04 | -0,152 ± 0,02 | -0,3389 ± 0,02 | -0,3142 ± 0,04 | -0,5344 ± 0,06 | -0,415 ± 0,05 |
| **Sum of neg. LFP** | mV | -17,56 ± 3,52 | -13,57 ± 2,03 | -33,31 ± 2,77 | -28,32 ± 3,36 | -51,07 ± 5,80 | -32,92 ± 3,58 |

**Mean (rank) difference**

| In. vs. W1 | In. vs. W2 | In. vs. W3 | In. vs. W4 | In. vs. W5 | W1 vs. W2 | W1 vs. W3 | W1 vs. W4 | W1 vs. W5 | W2 vs. W3 | W2 vs. W4 | W2 vs. W5 | W3 vs. W4 | W3 vs. W5 | W4 vs. W5 |
|---|---|---|---|---|---|---|---|---|---|---|---|---|---|---|
| -17,68 (ns) | -33,74 (**) | -49,6 (****) | -60,52 (**** | -46,59 (**** | -16,06 (ns) | -31,93 (*) | -42,84 (**** | -28,92 (*) | -15,87 (ns) | -26,78 (ns) | -12,86 (ns) | -10,91 (ns) | 3,01 (ns) | 13,92 (ns) |
| -0,04 (ns) | 0,03 (ns) | 0,04 (ns) | 0,08 (ns) | 0,01 (ns) | 0,07 (ns) | 0,09 (ns) | 0,12 (ns) | 0,06 (ns) | 0,02 (ns) | 0,05 (ns) | -0,01 (ns) | 0,03 (ns) | -0,03 (ns) | -0,06 (ns) |
| -37,88 (***) | -42,27 (***) | -27,82 (*) | -24,96 (ns) | -21,47 (ns) | -4,39 (ns) | 10,06 (ns) | 12,92 (ns) | 16,41 (ns) | 14,45 (ns) | 17,31 (ns) | 20,8 (ns) | 2,86 (ns) | 6,34 (ns) | 3,49 (ns) |
| -7,77 (ns) | -41,15 (***) | -34,87 (**) | -47,69 (**** | -34,64 (**) | -33,39 (*) | -27,10 (ns) | -39,92 (***) | -26,88 (ns) | 6,29 (ns) | -6,53 (ns) | 6,51 (ns) | -12,82 (ns) | 0,22 (ns) | 13,04 (ns) |
| -10,82 (ns) | -45,48 (**** | -36,11 (**) | -36,11 (**) | -35,87 (**) | -34,66 (**) | -25,28 (ns) | -25,28 (ns) | -25,05 (ns) | 9,37 (ns) | 9,37 (ns) | 9,61 (ns) | 0,0 (ns) | 0,24 (ns) | 0,24 (ns) |
| -18,21 (ns) | -42,90 (***) | -27,84 (*) | -33,61 (**) | -9,17 (ns) | -24,69 (ns) | -9,63 (ns) | -15,41 (ns) | 9,04 (ns) | 15,06 (ns) | 9,29 (ns) | 33,73 (*) | -5,77 (ns) | 18,67 (ns) | 24,44 (ns) |
| -11,82 (ns) | 21,0 (ns) | 17,43 (ns) | 37,58 (***) | 26,98 (ns) | 32,82 (*) | 29,25 (*) | 49,4 (****) | 38,8 (***) | -3,56 (ns) | 16,58 (ns) | 5,98 (ns) | 20,15 (ns) | 9,55 (ns) | -10,6 (ns) |
| -9,88 (ns) | 26,55 (ns) | 20,67 (ns) | 41,1 (***) | 23,18 (ns) | 36,43 (**) | 30,55 (*) | 50,98 (****) | 33,07 (**) | -5,88 (ns) | 14,55 (ns) | -3,36 (ns) | 20,43 (ns) | 2,51 (ns) | -17,91 (ns) |

Supplement, Table 2. **Values of MEA recordings.** All values given for every time period with corresponding unit, n-numbers and standard error of the mean. Each time period statistically compared with each other period and mean rank difference given with significance levels. Significance levels: ns = not significant, **** = p<0.0001, *** = p<0.001, ** = p<0.01, * = p<0.05

**RNA expression**

| Parameter | Unit | Initial (n=7) | Week 1 (n=7) | Week 2 (n=6) | Week 3 (n=8) | Week 4 (n=6) | |
|---|---|---|---|---|---|---|---|
| **Synaptophysin** | ddCt | 1 ± 0,11 | 2,033 ± 0,39 | 2,826 ± 1,02 | 3,311 ± 0,80 | 2,152 ± 0,29 | |
| **Parvalbumin** | ddCt | 1 ± 0,20 | 6,69 ± 0,84 | 15,63 ± 2,18 | 19,44 ± 2,77 | 12,87 ± 2,86 | |
| **HCN1** | ddCt | 1 ± 0,30 | 2,414 ± 0,35 | 2,826 ± 1,02 | 3,311 ± 0,80 | 2,152 ± 0,29 | - |
| **SCN1A** | ddCt | 1 ± 0,09 | 1,95 ± 0,13 | 2,121 ± 0,29 | 2,295 ± 0,33 | 1,826 ± 0,24 | - |
| **SCN2A** | ddCt | 1 ± 0,40 | 1,326 ± 0,40 | 0,7091 ± 0,24 | 0,9126 ± 0,23 | 0,6559 ± 0,12 | - |
| **SCN8A** | ddCt | 1 ± 0,09 | 1,606 ± 0,20 | 2,117 ± 0,10 | 2,118 ± 0,16 | 1,753 ± 0,21 | - |

**Mean (rank) difference**

| In. vs. W1 | In. vs. W2 | In. vs. W3 | In. vs. W4 | In. vs. W5 | W1 vs. W2 | W1 vs. W3 | W1 vs. W4 | W1 vs. W5 | W2 vs. W3 | W2 vs. W4 | W2 vs. W5 | W3 vs. W4 | W3 vs. W5 | W4 vs. W5 |
|---|---|---|---|---|---|---|---|---|---|---|---|---|---|---|
| -0,19 (ns) | -0,42 (ns) | -0,15 (ns) | 0,16 (ns) | - | -0,23 (ns) | 0,04 (ns) | 0,34 (ns) | - | 0,27 (ns) | 0,57 (ns) | - | 0,30 (ns) | - | - |
| -5,69 (ns) | -14,63 (***) | -18,44 (**** | -11,87 (**) | - | -8,94 (*) | -12,75 (***) | -6,18 (ns) | - | -3,81 (ns) | 2,76 (ns) | - | 6,57 (ns) | - | - |
| -1,41 (ns) | -1,83 (ns) | -2,31 (ns) | -1,15 (ns) | - | -0,41 (ns) | -0,90 (ns) | -0,26 (ns) | - | -0,48 (ns) | 0,68 (ns) | - | 1,16 (ns) | - | - |
| -0,95 (ns) | -1,12 (*) | -1,30 (**) | -0,83 (ns) | - | -0,17 (ns) | -0,35 (ns) | 0,12 (ns) | - | -0,17 (ns) | 0,30 (ns) | - | 0,47 (ns) | - | - |
| -0,33 (ns) | 0,29 (ns) | 0,09 (ns) | 0,34 (ns) | - | 0,62 (ns) | 0,41 (ns) | 0,67 (ns) | - | -0,20 (ns) | 0,05 (ns) | - | 0,26 (ns) | - | - |
| -0,61 (ns) | -1,12 (***) | -1,12 (****) | -0,75 (*) | - | -0,51 (ns) | -0,51 (ns) | -0,15 (ns) | - | -0,01 (ns) | 0,37 (ns) | - | 0,37 (ns) | - | - |

**Parvalbumin staining**

| Parameter | Unit | Initial (n=4) | Week 1 (n=4) | Week 2 (n=4) | Week 3 (n=4) | | |
|---|---|---|---|---|---|---|---|
| **Pvalb expression** | % | 0,175 ± 0,11 | 6,9 ± 0,54 | 11,1 ± 0,60 | 7,8 ± 1,63 | - | - |

**Mean (rank) difference**

| In. vs. W1 | In. vs. W2 | In. vs. W3 | In. vs. W4 | In. vs. W5 | W1 vs. W2 | W1 vs. W3 | W1 vs. W4 | W1 vs. W5 | W2 vs. W3 | W2 vs. W4 | W2 vs. W5 | W3 vs. W4 | W3 vs. W5 | W4 vs. W5 |
|---|---|---|---|---|---|---|---|---|---|---|---|---|---|---|
| -6,73 (**) | -10,93 (**** | -7,63 (***) | - | - | -4,2 (*) | -0,9 (ns) | - | - | 3,3 (ns) | - | - | - | - | - |

Supplement, Table 3. **Values of RT-PCR and parvalbumin staining.** All values given for every time period with corresponding unit, n-numbers and standard error of the mean. Each time period statistically compared with each other period and mean rank difference given with significance levels. Significance levels: ns = not significant, **** = p<0.0001, *** = p<0.001, ** = p<0.01, * = p<0.05
